# One-carbon Pathways and Methylation Potential in Glutamatergic Neurons Regulate Behavioral Alcohol Responses

**DOI:** 10.1101/2022.11.14.516474

**Authors:** Daniel Lathen, Collin Merrill, Gregory S. Ducker, Aylin R. Rodan, Adrian Rothenfluh

**Affiliations:** University of Utah, Huntsman Mental Health Institute, Dept. of Psychiatry; Interdepartmental Neuroscience Ph.D. Program; University of Utah, Dept. of Biochemistry; University of Utah, Molecular Medicine Program; University of Utah, Dept. of Internal Medicine, Division of Nephrology; University of Utah, Dept. of Human Genetics; University of Utah, Dept. of Neurobiology

## Abstract

Despite the enormous harms of alcohol use disorders (AUDs), many mechanisms, as well as effective prevention or treatment strategies remain elusive. Genetic factors dictate a majority of AUD risk. These risk factors can manifest as reduced naïve sensitivity to alcohol’s intoxicating effects and increased functional tolerance, i.e., brain-mediated decreases in sensitivity upon repeat exposure. The underlying neurobiology of how AUD-associated genes alter these endophenotypes remains poorly understood. Genes implicated in AUDs include epigenetic modifiers, such as histone demethylases, including *Kdm3*. We previously showed that whole-body and neuronal *Kdm3* strongly affect ethanol sensitivity and tolerance in *Drosophila*. Here, we investigate the mechanisms of these effects, and, by extension, mechanisms of sensitivity and tolerance. RNA-seq and pathway analysis on *Kdm3*^*KO*^ flies revealed disproportionate upregulation of genes involved in amino acid metabolism, including 1-carbon pathways. We show that acute amino acid feeding modulates sensitivity and tolerance in a *Kdm3*-dependent manner. Global manipulation of 1-carbon genes, especially glycine *N*-methyltransferase (*Gnmt*), glycine decarboxylase (*Gldc*), and sarcosine dehydrogenase (*Sardh*), alters alcohol sensitivity and tolerance. These changes in alcohol responses are likely mediated by global glycine levels (a substrate of these enzymes) rather than by 1-carbon input. Conversely, neuronal manipulations of 1-carbon pathways change alcohol sensitivity and tolerance in a pattern that suggests a mechanism through *S*-adenosyl methionine (SAM), a 1-carbon metabolite that is the universal methyl donor required for epigenetic methylation. Increasing SAM production specifically in glutamatergic neurons increases sensitivity and tolerance. Together, these findings reveal distinct mechanisms affecting alcohol sensitivity and tolerance globally (via glycine) and neuronally (via SAM), thus revealing an important and complex role of 1-carbon metabolism in mediating AUD phenotypes.

## INTRODUCTION

Alcohol use disorders (AUDs) exact an immense toll on individuals, families, and society. Genetic factors determine up to 60% of an individual’s risk of developing problematic alcohol habits (Goldman, Oroszi, & Ducci, 2005). To better identify genetic factors, alcohol addiction can be broken down into discrete endophenotypes, including naïve sensitivity and rapid functional tolerance (i.e., brain-mediated decreases in sensitivity measured after all EtOH from initial exposure has completely metabolized). The degree to which an individual displays reduced naïve sensitivity to EtOH and develops rapid functional tolerance suggests their propensity for developing AUDs (Atkinson, 2009; Mayfield, Harris, & Schuckit, 2008; Schuckit, 2009). Yet, the genetic, neuronal, and molecular mechanisms behind these two EtOH responses are not well understood (Goldman et al., 2005; Park, Ghezzi, Wijesekera, & Atkinson, 2017; Rodan & Rothenfluh, 2010; Scholz, Ramond, Singh, & Heberlein, 2000).

The genetic amenability of *Drosophila melanogaster* makes it an excellent model system for discovering conserved genes and elucidating mechanisms. Vinegar flies exhibit strong face and mechanistic validity as models for EtOH abuse (Gonzalez et al., 2018; Narayanan & Rothenfluh, 2016; Ojelade, Jia, et al., 2015; Park et al., 2017; Robinson & Atkinson, 2013; Rodan & Rothenfluh, 2010). Like humans, flies become hyperactive upon exposure to low doses of EtOH but sedate at high doses (Rodan, Kiger, & Heberlein, 2002). Further, *Drosophila* readily develop rapid functional tolerance to EtOH (Berger, Heberlein, & Moore, 2004; Scholz et al., 2000). This tolerance is not due to altered pharmokinetics (Scholz et al., 2000) and is similar to rapid ethanol tolerance in rodents, which is a proxy of AUD-associated chronic tolerance (Kalant, 1998; Khanna, Kalant, Shah, & Weiner, 1991; Lê & Kiianmaa, 1988; Rustay & Crabbe, 2004). *Drosophila* also exhibit strong predictive validity, as exemplified by unbiased alcohol studies implicating *Drosophila* orthologs of genes that are subsequently implicated in human alcoholism (Gonzalez et al., 2018).

Recent studies using *Drosophila* and other model systems support an important role of histone demethylases (HDMs) in AUD-associated behaviors (T. D. Berkel & Pandey, 2017; Ramirez-Roman, Billini, & Ghezzi, 2018; Shukla et al., 2008). Exposure to alcohol and other drugs of abuse alters gene expression by changing cells’ epigenetic landscapes (T. D. Berkel & Pandey, 2017; Shukla et al., 2008). In addition, epigenetic modifiers such as histone methyltransferases (HMTs) and HDMs affect EtOH phenotypes, likely by influencing transcriptional control (Barbier et al., 2017; T. D. Berkel & Pandey, 2017; T. D. M. Berkel, Zhang, Teppen, Sakharkar, & Pandey, 2019; Maze et al., 2010; Pinzon et al., 2017; Ponomarev, 2013; Qiang, Denny, Lieu, Carreon, & Li, 2011; Zakhari, 2013). We previously showed that among all *Drosophila* HDM orthologs, loss of lysine-specific histone demethylase 3 (*Kdm3)* strongly alters alcohol phenotypes, increasing naïve sensitivity and decreasing tolerance (Pinzon et al., 2017). However, as is true for most AUD-associated genes, how *Kdm3* produces these responses is unknown.

HMTs require *S*-adenosyl methionine (SAM), the universal methyl donor for methyltransferases. SAM is a key output of the methionine cycle and the folate cycle, which together form the core of 1-carbon (1-C) metabolism. Recent evidence suggests that SAM, 1-C enzymes, and HDMs influence histone methylation (Friso, Udali, De Santis, & Choi, 2017; Liu, Barnes, & Pile, 2015; Mentch & Locasale, 2016; Mentch et al., 2015; Serefidou, Venkatasubramani, & Imhof, 2019).

We investigated the mechanisms through which histone modifiers like *Kdm3* modulate AUD phenotypes. Here, we show that *Kdm3* is linked to 1-C metabolism and that 1-C metabolites and enzymes alter ethanol sensitivity and tolerance. Globally, glycine and enzymes involved in glycine metabolism alter alcohol phenotypes, whereas in neurons, particularly glutamatergic neurons, SAM mediates alcohol sedation sensitivity and tolerance.

## RESULTS

### Genes associated with 1-carbon metabolism are upregulated in *Kdm3*^*KO*^ flies

As an epigenetic modifier, we hypothesized that *Kdm3* loss would result in substantial changes in gene expression, which might in turn affect alcohol responses. Therefore, we first asked how knocking out *Kdm3* altered gene expression. RNA-seq analysis of *Kdm3*^*KO*^ fly heads revealed 359 downregulated genes and 457 upregulated genes (Fig. 1A). Gene ontology and pathway enrichment analysis revealed that pathways associated with amino acid metabolism were enriched for upregulated genes (Fig. 1B; *p* = 2.99×10^−14^, Benjamini-Hochberg adjusted). Many of these genes are components of 1-C pathways, including glycine N-methyltransferase (*Gnmt*), serine hydroxymethyl transferase (*Shmt*), and glycine dehydrogenase (*Gldc*, a.k.a. *CG3999* in flies) (Fig. 1A). Subsequent RT-qPCR analysis confirmed elevated transcription of *Gnmt* and *Gldc*, but not of *Shmt* (Fig. 1C).

**Fig. 1.**
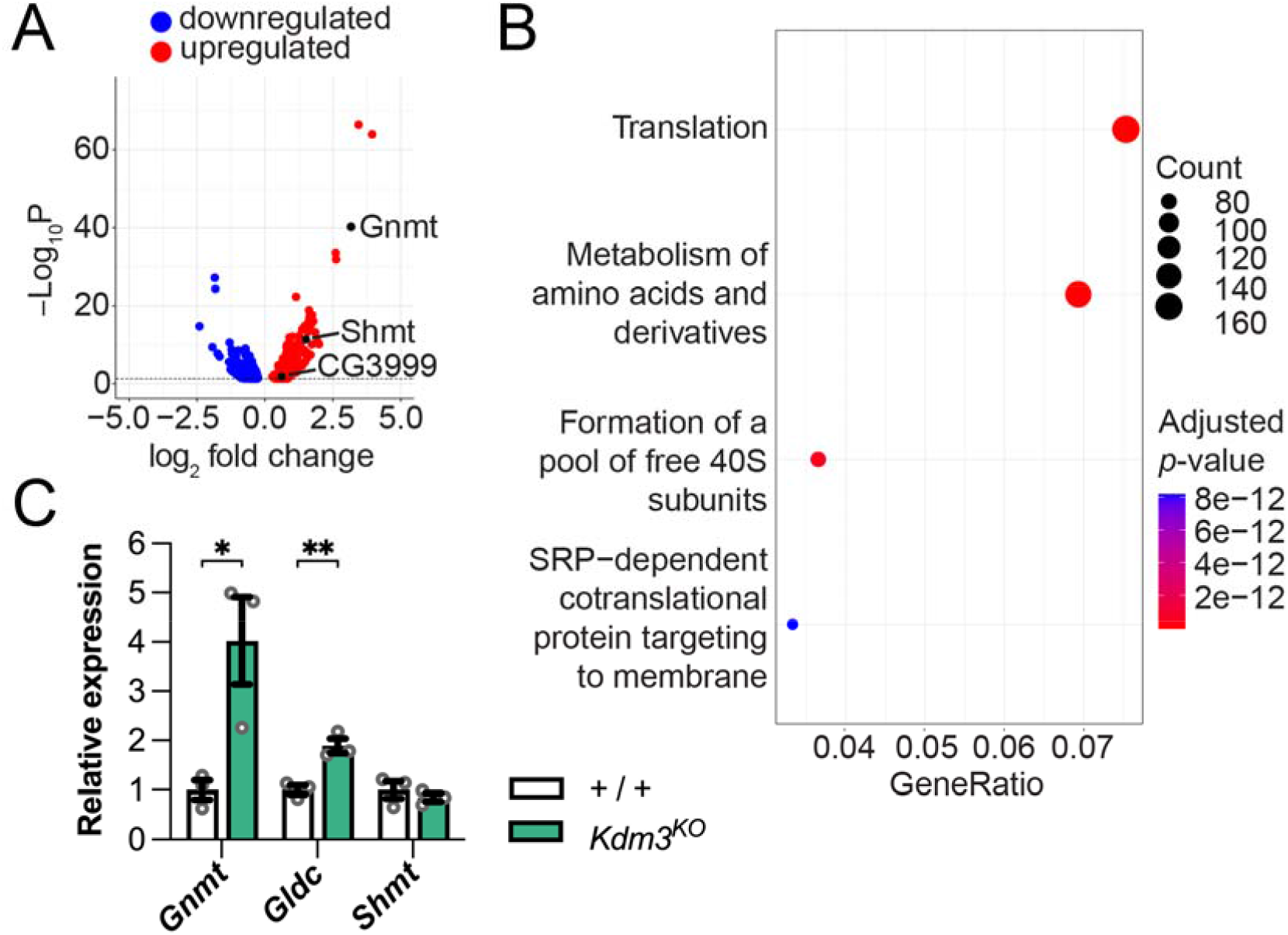
Genes associated with 1-C metabolism are upregulated in *Kdm3*^*KO*^ flies. (A) Significantly upregulated (red) or downregulated (blue) transcripts in *Kdm3*^*KO*^ fly heads. (B) Gene ontology (GO) and enriched pathway analysis of genes upregulated in *Kdm3*^*KO*^ flies showing the top four overrepresented pathway terms. Gene ratio indicates the ratio of genes in the dataset to all genes associated with GO pathways. (C) RT-qPCR confirming *Gnmt* and *Gldc* upregulation in *Kdm3*^*KO*^ fly heads. Gene expression was normalized to *Rpl32* expression. Statistical differences were analyzed using Student’s *t-*tests in (C) and in all subsequent two-group comparisons. The data in (C) and in all subsequent graphs represent the mean ± SEM, with \**p* < 0.05; \*\**p* < 0.01; \*\*\**p* < 0.001. *n* = 3.

### Glycine decreases alcohol sensitivity and tolerance in a *Kdm3*-dependent manner

Because *Kdm3* knockout upregulates genes involved in amino acid metabolism and 1-C pathways, we hypothesized that changes in amino acids may play a role in *Kdm3-* mediated alcohol phenotypes. To test this hypothesis, we fed adult flies casamino acids for three days before testing alcohol sensitivity and tolerance. Casamino acids are a mixture of all primary amino acids except tryptophan. In control flies, casamino acid feeding dose-dependently decreased alcohol sensitivity without affecting tolerance (Fig. 2A). To determine whether altered amino acid metabolism affects alcohol phenotypes via *Kdm3*, we performed equivalent experiments using *Kdm3*^*KO*^ flies. *Kdm3* knockout abolished the casamino acid-induced sensitivity to alcohol (Fig. 2B). To home in on which amino acids underlie the observed resistance to alcohol sedation, we fed flies glycine, which is a substrate for *Gnmt* and *Gldc*. Glycine feeding caused decreased sensitivity and tolerance in control flies (Fig. 2C). Again, these effects were abolished in *Kdm3*^*KO*^ flies (Fig. 2D). The alcohol sedation and tolerance differences between controls and *Kdm3*^*KO*^ flies were not due to lower amino acid consumption by *Kdm3*^*KO*^ flies compared to controls (Fig. S1). Together, these results suggest that glycine is involved in alcohol-induced sedation sensitivity and tolerance via a *Kdm3*-dependent mechanism.

**Fig. 2.**
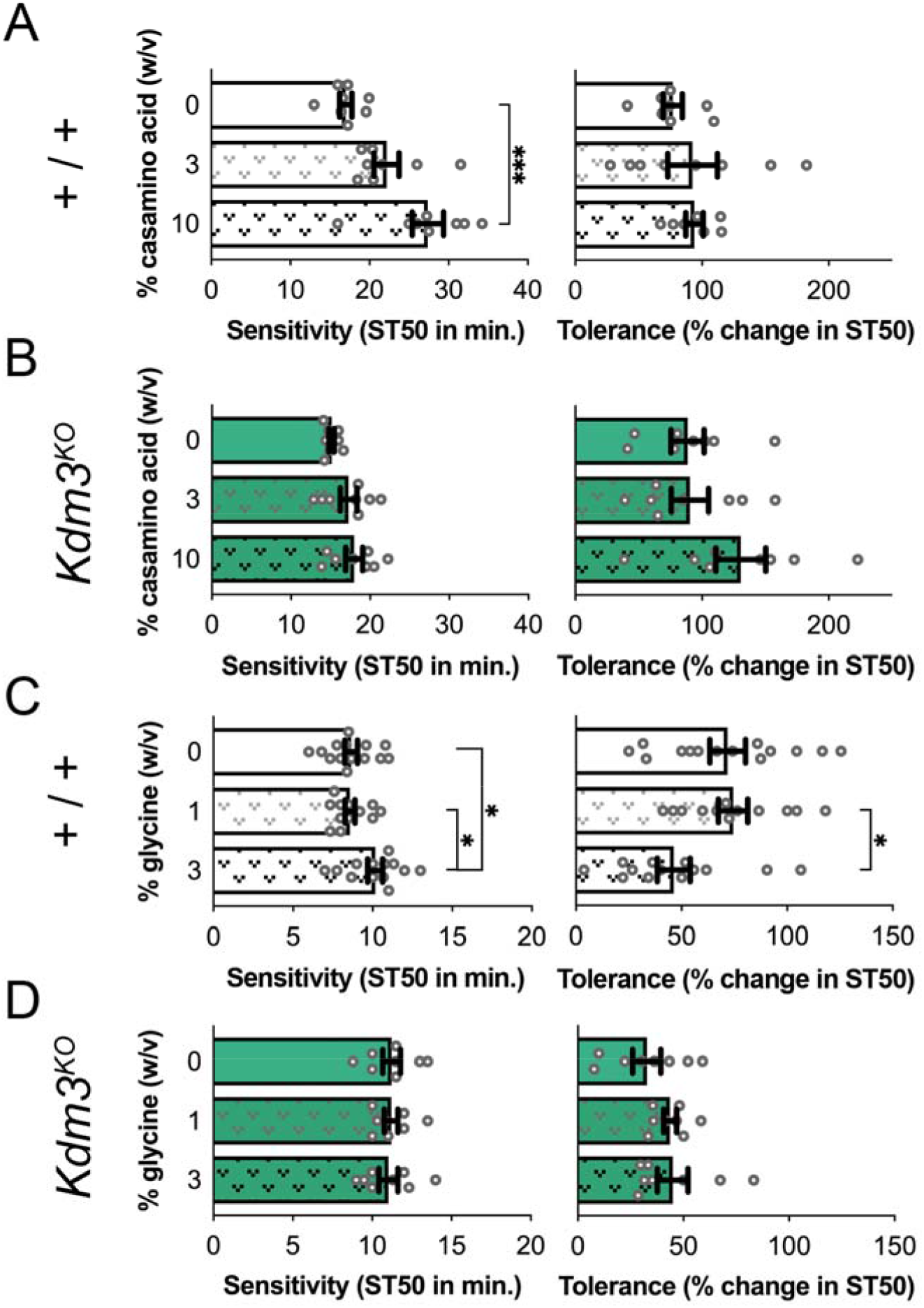
Glycine feeding decreases sensitivity and tolerance in a *Kdm3*-dependent manner. (A-B) Casamino acid consumption for three days decreased sensitivity in control flies (A) but not in *Kdm3*^*KO*^ flies (B). Sensitivity was measured as the time at which 50% of the flies in each vial were sedated (ST50). A higher initial ST50 represents reduced sensitivity. Tolerance was measured as the percent change between the ST50 at the second exposure and the initial ST50. (C-D) Glycine consumption for three days decreases sensitivity and tolerance in control flies (C) but not in *Kdm3*^*KO*^ flies (D). Statistical differences were analyzed using one-way ANOVA followed by Tukey post-hoc tests, as well as in all subsequent three-group comparisons. *n* ≥ 7. Each dot in (A-B) represents 20 flies. Each dot in (C-D) and all other EtOH experiments represents 10 flies.

### Increased folate cycle activity is not a primary driver of sensitivity and tolerance

Through *Gldc*, glycine can be a source of methyl groups fueling the 1-C methionine cycle (Fig. 3A). Glycine could decrease sensitivity and tolerance by increasing folate cycle activity. In this case, *Gldc* loss would decrease methyl group input and cause the opposite phenotype as glycine feeding. We tested this hypothesis using a *Gldc* null mutant, *Gldc*^*Mi*^, which is an intronic Minos-mediated integration cassette (MiMIC) gene trap insertion (Venken et al., 2011). qPCR using probes spanning the MiMIC-containing intron yielded no amplification, indicating aberrant transcription. Mutating one or both copies of *Gldc* led to decreased alcohol sensitivity and tolerance instead of the expected opposite phenotype, suggesting that input into the folate cycle does not underlie the glycine-induced phenotypes we observed (Fig. 3B-C). *Gldc* loss increases glycine levels (Leung et al., 2017; Pai et al., 2015), so we wished to determine if increased glycine levels themselves caused the glycine feeding and *Gldc*^*Mi*^ results. To that end, we tested whether *Gldc* mutation potentiated glycine-induced phenotypes. Interestingly, *Gldc*^*Mi*^ flies died when fed 1% or 3% glycine, suggesting that *Gldc* plays a critical role in glycine breakdown. Indeed, *Gldc* loss induces glycine-dependent toxicity in mouse brain (Kim et al., 2015), and elevated glycine levels cause cessation of feeding and growth in flies (Zinke, Kirchner, Chao, Tetzlaff, & Pankratz, 1999). When previously fed 3% glycine, control flies showed decreased alcohol sensitivity and tolerance (Fig. 2C). When fed only 0.3% glycine, control flies showed no changes in alcohol sensitivity or tolerance (Fig. 3D). As expected, *Gldc*^*Mi*^ flies again showed decreased sensitivity to alcohol and decreased tolerance compared to control flies. In contrast with controls, feeding *Gldc*^*Mi*^ flies with 0.3% glycine did not induce a sedation phenotype, but tolerance was decreased (Fig. 3D). We detected significant main effects of genotype for both sensitivity and tolerance and a significant genotype x glycine feeding interaction for tolerance. This suggests that *Gldc* loss exacerbates rather than eliminates the effects of glycine. Therefore, glycine levels, rather than input into the folate cycle, mediate our glycine-feeding and *Gldc* phenotypes. Also consistent with this hypothesis, flies containing a putative global *Shmt* mutation that disrupts serine-dependent folate cycle input did not show altered alcohol sensitivity or tolerance (Fig. 3E). Finally, to confirm that the folate cycle is not involved in glycine-induced resistance to alcohol, we fed flies with formate for three days. Formate augments methylene-tetrahydrofolate (CH_2_-THF) to act as an alternate carbon source parallel to *Gldc*-dependent CH_2_-THF synthesis from glycine (Brosnan & Brosnan, 2016). We found no significant effects from supplemental formate (Fig. 3F). Together, our results suggest that the folate cycle is not involved in glycine-mediated decreases in alcohol sensitivity and tolerance, but rather that glycine levels themselves are responsible. Further supporting this hypothesis, glycine may fail to produce phenotypes in *Kdm3*^*KO*^ flies (Fig. 2D) because upregulation of *Gldc* and *Gnmt* (Fig. 1A,C) enhances glycine catabolism and prevents glycine buildup, suggesting that glycine rather than its metabolic products creates the alcohol phenotypes.

**Fig. 3.**
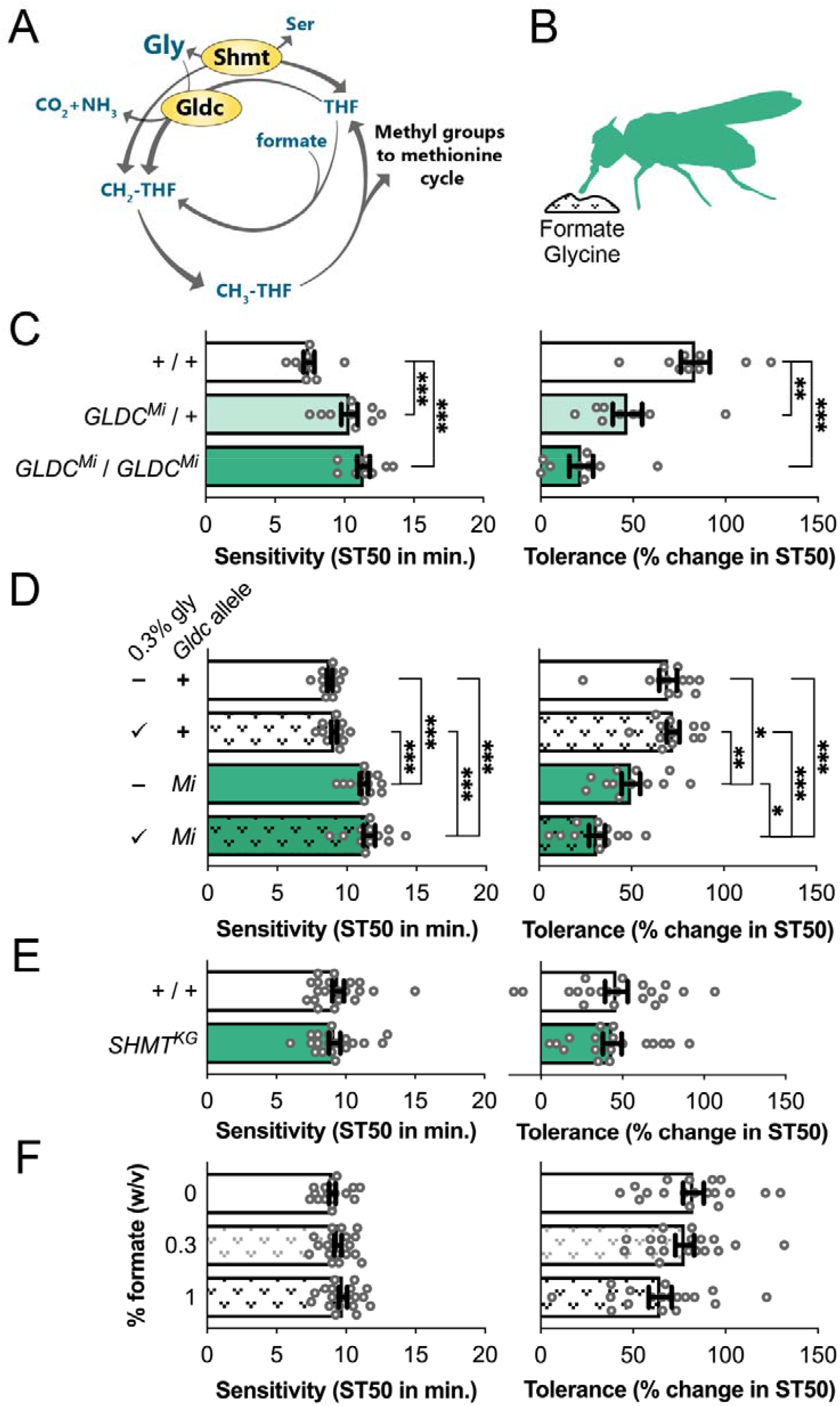
Increased folate cycle activity is not a primary driver of alcohol sensitivity or tolerance. (A) Schematic of the folate cycle. (B) Schematic of manipulation locations. Green represents whole-body manipulations. (C) Heterozygous or homozygous *Gldc* null mutation decreases alcohol sensitivity and tolerance. (D) Low glycine feeding had no effect on sedation sensitivity in control or *Gldc* mutants but had a synergistic effect on tolerance in *Gldc* mutants. (*Sensitivity:* glycine main effect, *p* = .236; genotype main effect, *p* < .001; interaction, *p* = .896. *Tolerance:* glycine main effect, *p* = .087; genotype main effect, *p* < .001; interaction, *p* = .022). Statistical differences were analyzed using two-way ANOVA followed by Tukey post-hoc tests, as well as in all subsequent two-factor comparisons. (E) *Shmt* knockout mutation does not affect alcohol sensitivity or tolerance. (F) Formate feeding for three days does not affect alcohol sensitivity or tolerance. *n* ≥ 9.

### *Gnmt* modulates glycine-induced changes to alcohol sensitivity and tolerance

Since glycine did not affect alcohol sensitivity and tolerance via its role in the folate cycle, we next examined its role in the methionine cycle. *Gnmt*, which is upregulated in *Kdm3*^*KO*^ flies, consumes glycine in the latter cycle (Fig. 4A). Using *Gnmt*^*Mi*^, a validated null mutation (Obata & Miura, 2015), we found that global *Gnmt* loss of either one or both alleles decreased sensitivity and tolerance (Fig. 4B-C). These data indicate haploinsufficiency of *Gnmt*, similar to *Gldc*, and sensitivity to total enzyme activity. We corroborated our whole-body *Gnmt*^*Mi*^ results with whole-body *Gnmt* knockdown using *tubulin84B*-*Gal4* driven *Gnmt*^*RNAi*^ (Fig. 4D). Notably, the *tubulin84B-Gal4* driver does not induce alcohol phenotypes (Supp. 2A). A second RNAi construct targeting a distinct region of *Gnmt* and validated by other groups (Obata et al., 2014; Obata & Miura, 2015) caused the same phenotype (Supp. 3A-B). Both RNAi knockdowns yielded the same result as *Gnmt* knockout and glycine feeding (i.e. decreased sensitivity and tolerance; Fig. 4D and Supp. 3A-B). Augmenting glycine may enhance *Gnmt* activity, so we again performed 2-way ANOVA to determine if *Gnmt* is necessary for glycine-induced phenotypes. Indeed, global *Gnmt*^*Mi*^ expression abolished sensitivity and tolerance phenotypes induced by glycine feeding in control flies (Fig. 4E; we detected a significant glycine main effect for sensitivity and a significant interaction for both sensitivity and tolerance). This reduced phenotype could indicate a ceiling effect. However, a similar possible ceiling effect from excess glycine in glycine-fed *Gldc* mutants caused lethality, whereas *Gnmt* mutants tolerated 3% gly feeding. Therefore, our result showing occluded glycine-induced reductions in sensitivity and tolerance could alternatively suggest that, unlike *Gldc, Gnmt* is required for the effects of glycine on alcohol sensitivity and tolerance. Thus, glycine may modify alcohol phenotypes via *Gnmt*-dependent buffering of SAM (Fig. 4A). To further disrupt this buffering capability, we used an RNAi construct (Obata et al., 2014; Obata & Miura, 2015) to globally knock down *Sardh*, an enzyme that synthesizes glycine and performs the inverse function of *Gnmt* (Fig. 4A). *Sardh* knockdown decreased alcohol sensitivity and tolerance (Fig. 4F) similar to *Gnmt* loss, despite opposite expected effects on glycine. Thus, the functionality of the *Gnmt*-*Sardh* cycle may predict alcohol phenotypes better than glycine levels alone.

**Fig. 4.**
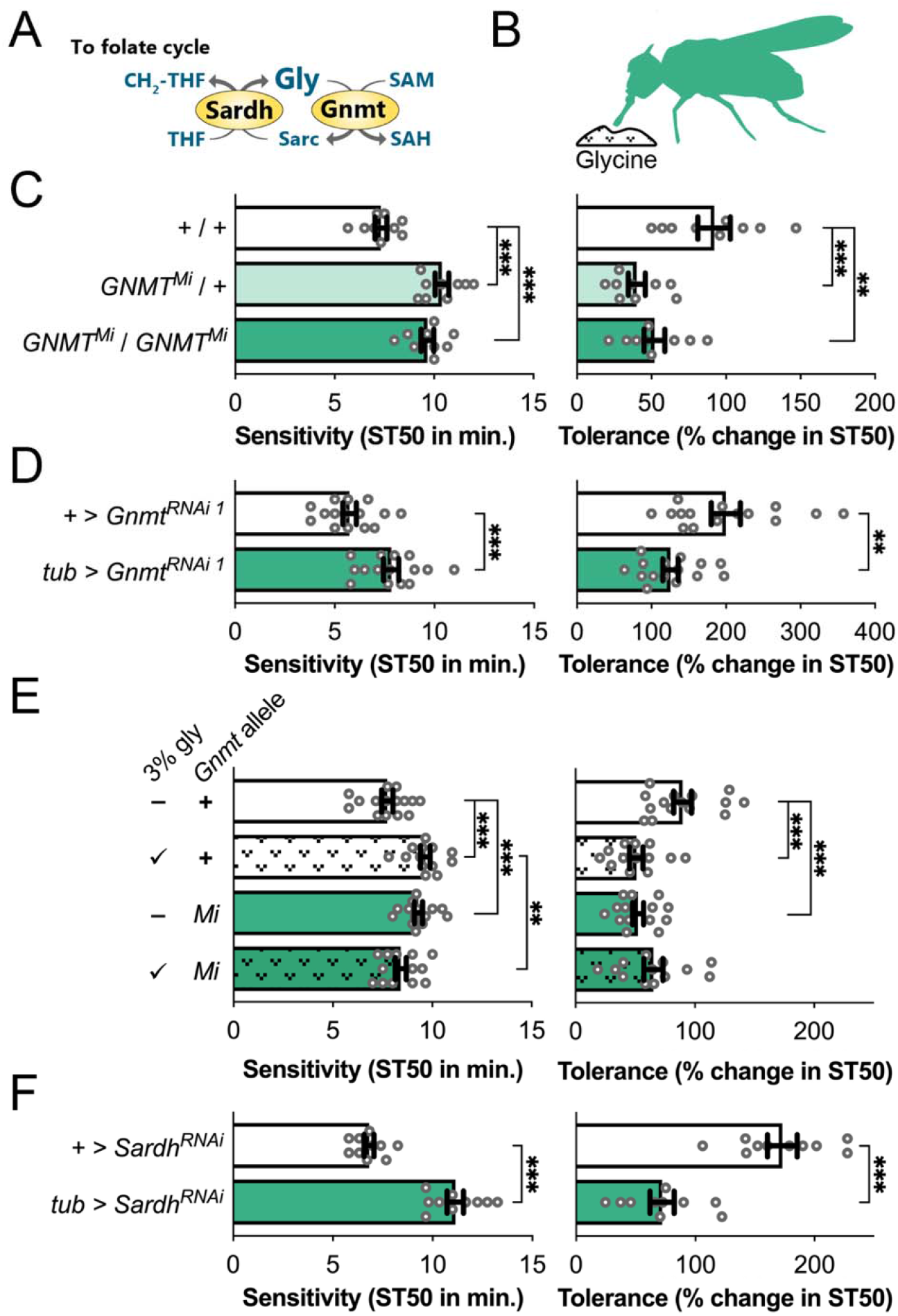
*Gnmt* modulates glycine-induced changes to alcohol sensitivity and tolerance. (A) Schematic depicting the relationships between *Gnmt, Sardh*, and glycine. (B) Color-coded schematic of manipulation locations. (C) Heterozygous or homozygous *Gnmt* knockout decreases alcohol sensitivity and tolerance. (D) Whole-body *Gnmt* knockdown driven by *Tubulin84B-Gal4* decreases alcohol sensitivity and increases tolerance. (E) Control flies fed 3% glycine show decreased sensitivity and tolerance to alcohol-induced sedation, while *Gnmt*-null mutants show no changes. Glycine feeding and *Gnmt* knockout show inhibitory interaction in both sensitivity and tolerance (*Sensitivity*: glycine main effect, *p* = .049; genotype main effect, *p* = .512; interaction, *p* < .001. *Tolerance*: glycine main effect, *p* = .055; genotype main effect, *p* = .087; interaction, *p* < .001). (F) Whole-body *Sardh* knockdown decreases alcohol sensitivity and tolerance. *n* ≥ 9.

### Neuronal *S*-adenosyl-methionine (SAM) increases sensitivity and tolerance

Loss of *Gnmt* or *Sardh* may both increase SAM levels by interrupting *Gnmt* buffering in the methionine cycle (Fig. 5A). Indeed, *Gnmt* mutants and *Sardh* mutants exhibit increased SAM levels (Kashio et al., 2016; Luka, Capdevila, Mato, & Wagner, 2006; Obata et al., 2014). Thus, we next tested the hypothesis that *Gnmt* acts via SAM to alter alcohol responses. Opposite phenotypes from *Gnmt* loss (i.e., higher SAM) and SAM reduction would support this hypothesis. However, when we fed flies cycloleucine, an inhibitor of SAM synthase (*SamS*) (Sufrin, Coulter, & Talalay, 1979), we observed decreased sensitivity and tolerance, similar to results using *Gnmt*^*Mi*^ flies (Fig. 5B-C). Since ethanol sensitivity and tolerance are neuronal phenomena (Robinson & Atkinson, 2013; Rodan et al., 2002; Scholz et al., 2000) and numerous genes are required in neurons for normal sensitivity and tolerance (Engel et al., 2016; Ojelade, Acevedo, Kalahasti, Rodan, & Rothenfluh, 2015; Ojelade, Jia, et al., 2015; Pinzon et al., 2017; B. R. Troutwine, Ghezzi, Pietrzykowski, & Atkinson, 2016), we next shifted our focus to neurons. We hypothesized that altering *SamS* in neurons is sufficient to alter alcohol responses. First, we verified that the pan-neuronal driver *elav-Gal4* alone did not cause alcohol phenotypes (Supp. 3B). Expressing *SamS*-RNAi neuronally was lethal in males but significantly reduced sensitivity and tolerance in females (Fig. 5D) (Obata & Miura, 2015). This result was consistent with cycloleucine feeding. Furthermore, *SamS* overexpression (Obata & Miura, 2015), which increases SAM levels, caused increased sensitivity and tolerance (Fig. 5E), suggesting that alcohol sensitivity is correlated with SamS levels. Next, we increased SAM levels by knocking down *Gnmt* in neurons using both RNAi constructs (Luka et al., 2006; Obata et al., 2014). Consistent with neuronal *SamS* overexpression, neuronal *Gnmt* knockdown caused increased alcohol sensitivity and tolerance (Fig. 5F and Supp. 3C). Though *Gnmt* is expressed at low levels in the brain and in neurons (Li et al., 2022), our results indicate that neuronal *Gnmt* plays an important role in alcohol sensitivity and tolerance. Taken together, these results suggest that neuronal SAM levels affect alcohol sensitivity and tolerance phenotypes.

**Fig. 5.**
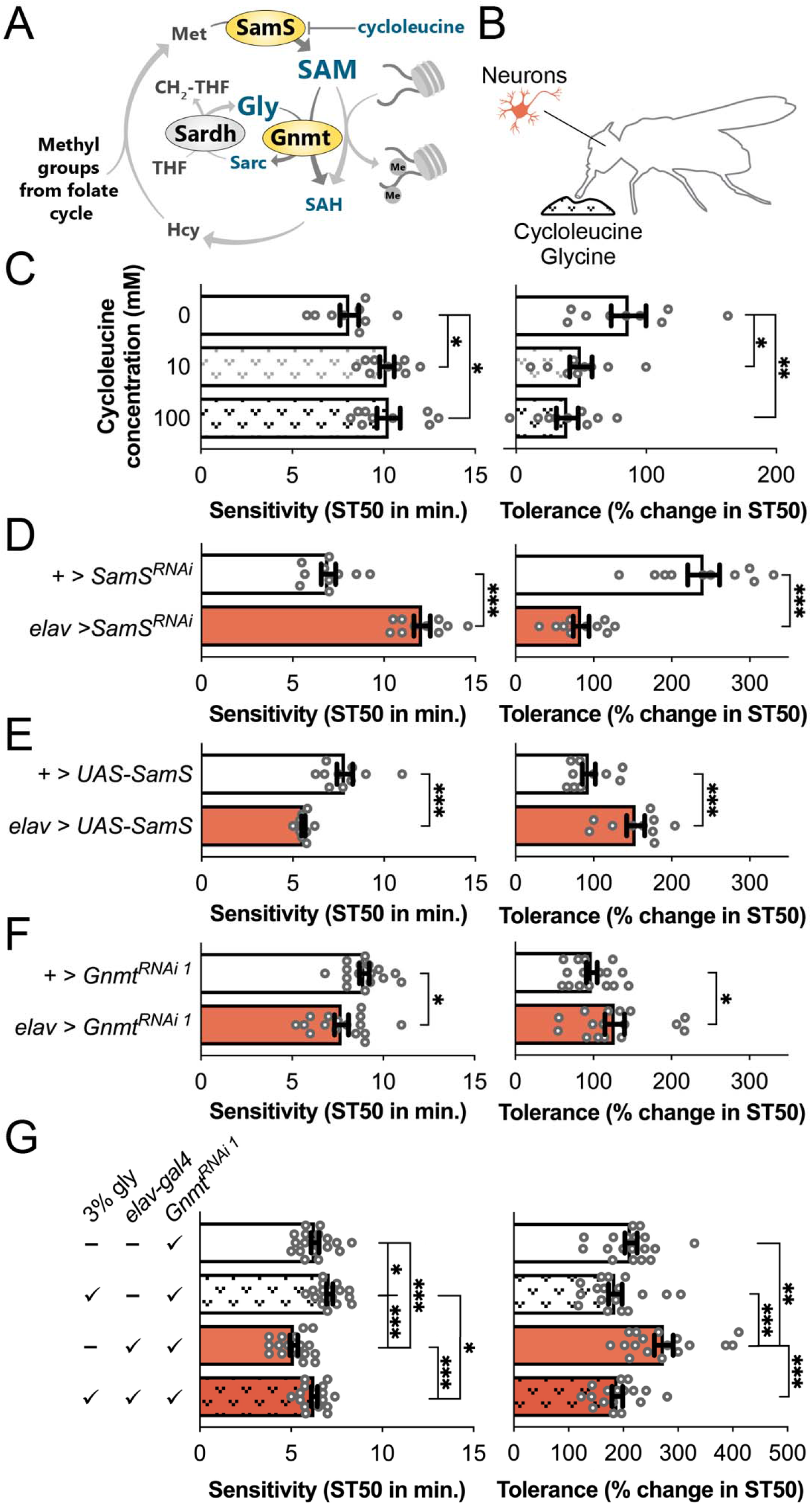
Neuronal *S*-adenosyl methionine (SAM) increases alcohol sensitivity and tolerance. (A) Schematic of the methionine cycle. (B) Color-coded schematic of manipulation locations. (C) Cycloleucine feeding for three days decreases alcohol sensitivity and tolerance. (D) Pan-neuronal *SamS* knockdown using *elav-Gal4* and *UAS-SamS*^*RNAi*^ substantially decreases sensitivity and tolerance, whereas pan-neuronal *SamS* overexpression shows the opposite effect. (E) Pan-neuronal *Gnmt* knockdown increases alcohol sensitivity and tolerance. (F) 2-way ANOVA with glycine feeding and neuronal *Gnmt* knockdown shows no sensitivity interaction and slight tolerance interaction. (*Sensitivity*: glycine main effect, *p* < .001; genotype main effect, *p* < .001; interaction, *p* = .406. *Tolerance*: glycine main effect, *p* < .001; genotype main effect, *p* = .016; interaction, *p* = .034). *n* ≥ 9.

Surprisingly, neuronal *Gnmt* knockdown caused a phenotype opposite to global *Gnmt* loss, suggesting that *Gnmt* has distinct mechanisms of action in neurons compared to the whole body (i.e., SAM acts in neurons while glycine acts in the body). Consistent with this hypothesis, glycine feeding and global *Gnmt* null mutation previously showed an interaction (Fig. 4E; green bars), but glycine feeding in neuronal *Gnmt*^*RNAi 1*^ knock down showed no interaction for the sensitivity phenotypes and only a subtle interaction on tolerance (Fig. 5G; orange bars). These data suggest that whole-body *Gnmt* loss affects the same pathways as glycine feeding, whereas neuronal *Gnmt* loss primarily affects pathways distinct from global glycine. The slight tolerance interaction may indicate that the global glycine effects dominate over the effects of neuronal *Gnmt*.

Glycine is used as an inhibitory neurotransmitter in the brain, so altered glycine metabolism could affect glycinergic neurotransmission to produce neuronal *Gnmt* knockdown phenotypes. Thus, we disrupted glycinergic neurotransmission by knocking down a glycine receptor gene (*Grd* (Frenkel et al., 2017)) and a glycine synaptic reuptake transporter (*GlyT*; Supp. 4A). Knocking down *Grd* in all neurons increased alcohol sedation and tolerance (Supp. 4B), while knocking down *GlyT* only increased tolerance (Supp. 4C). If elevated glycine or glycinergic signaling explained neuronal *Gnmt* phenotypes, we would expect opposing results from *Gnmt* loss (and expected subsequent glycine increases) and interruption of glycinergic signaling, but this was not the case. These data suggest that SAM, rather than glycine or glycinergic neurotransmission, mediates neuronal sensitivity and tolerance to alcohol.

Neuronal SAM levels could be regulated via mechanisms external to neurons, such as input from the fat body or by glia. The fat body regulates metabolism and energy storage, similar to the human liver and adipocytes (Rizki & Rizki, 1978). Further, *Gnmt, Sardh, Shmt*, and *Gldc* are highly expressed in the fat body (Li et al., 2022). Additionally, *Gnmt* expressed in the fat body is central to SAM regulation in flies (Obata et al., 2014). Glia are critical for regulating synaptic levels of neurotransmitters and other molecules (Y. Kim, Park, & Choi, 2019). Therefore, we tested if the fat body or glia might control our observed global phenotypes by knocking down *Gnmt* in these structures using RNAi (Supp. 5A). However, our results phenocopied *Gnmt* loss in neurons rather than *Gnmt* loss in whole flies (Supp. 5B-C). Though the identity of the non-neuronal tissue determining the global *Gnmt* phenotype remains to be determined, our experiments indicate that neuronal SAM modulates alcohol sensitivity and tolerance.

### SAM in glutamatergic neurons modulates sensitivity and tolerance

Alcohol-induced sedation can occur via dysregulation of the homeostatic balance between competing excitatory and inhibitory neuron activity (Ghezzi, Li, Lew, Wijesekera, & Atkinson, 2017). Glycine may influence the former via its role as required co-agonist of NMDA-type glutamate receptors (NMDAR). Further, although no glycinergic neuron-specific *Gal4* driver exists, glycine is often co-released with GABA at inhibitory synapses, and both neurotransmitters rely on the vesicular transporter VGAT (Aubrey & Supplisson, 2018). Thus, we knocked down *Gnmt* in excitatory glutamatergic and inhibitory VGAT-expressing neurons (Fig. 6A). *Gnmt* knockdown in inhibitory neurons did not affect sedation or tolerance (Fig. 6B). In contrast, *Gnmt* knockdown in glutamatergic neurons increased alcohol sensitivity and tolerance, similar to pan-neuronal *Gnmt* knockdown (Fig. 6C). These results suggest that SAM activity in glutamatergic neurons regulates alcohol phenotypes.

**Fig. 6.**
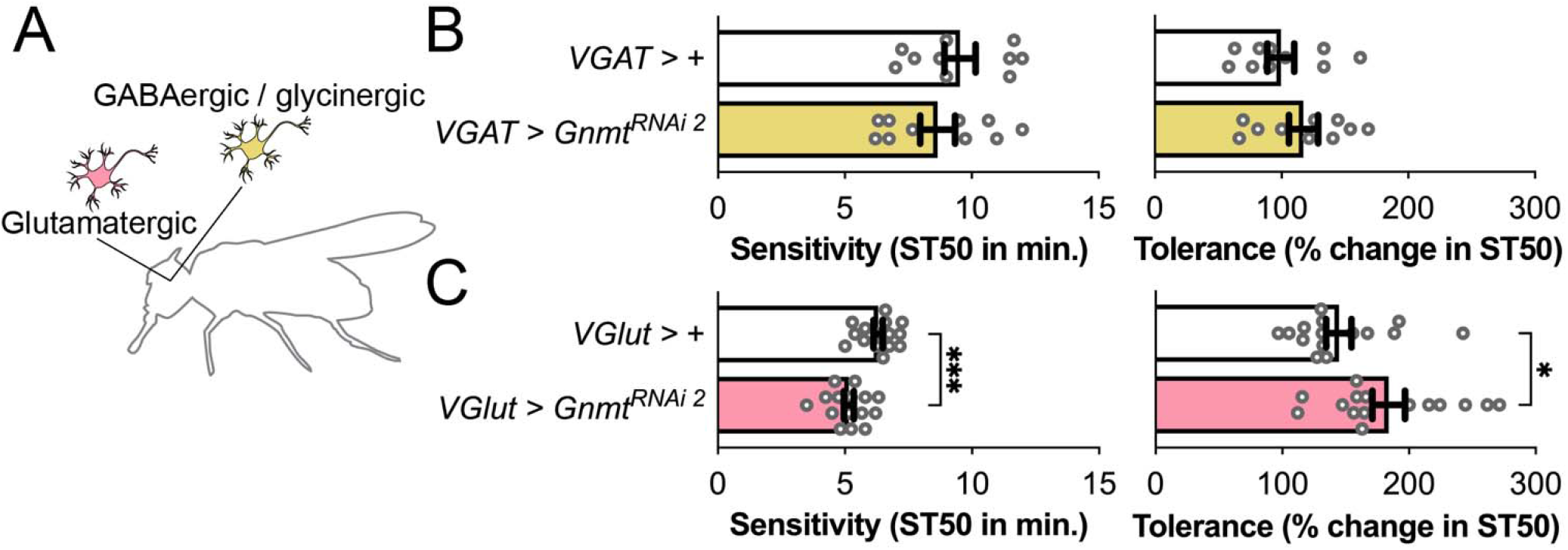
*S*-adenosyl methionine (SAM) in glutamatergic neurons modulates alcohol sensitivity and tolerance. (A) Color-coded schematic of manipulated neurons. (B) *Gnmt* loss in inhibitory neurons using *VGAT-Gal4*, a GABAergic/glycinergic neuron-specific driver, did not affect alcohol sedation and tolerance, whereas *Gnmt* loss in glutamatergic neurons using *vGlut-Gal4*, a glutamatergic neuron-specific driver, increased alcohol sensitivity and tolerance. *n* ≥ 10.

## DISCUSSION

The present study investigates *Kdm3*-dependent mechanisms of alcohol sedation sensitivity and tolerance. In so doing, we have elucidated a role of amino acids and 1-C metabolism in modulating AUD phenotypes, culminating in our finding that SAM levels in glutamatergic neurons regulate alcohol sensitivity and tolerance phenotypes (Fig. 7). We show that loss of *Kdm3* upregulates expression of 1-C enzymes. These genes in turn affect alcohol phenotypes, consistent with *Kdm3* playing a role in regulating alcohol behaviors (Pinzon et al., 2017). Many other studies indicate that *Kdm3* plays a critical role in alcohol abuse (Mulligan et al., 2006; Ponomarev, Wang, Zhang, Harris, & Mayfield, 2012; Qiang et al., 2011; Subbanna et al., 2013). An outstanding question is how *Kdm3* loss changes expression of these genes. As a histone modifier, *Kdm3* may directly regulate these genes by demethylating their associated histones. Alternatively, *Kdm3* loss may affect expression of 1-C genes by altering methyl group availability and necessitating compensatory homeostatic responses. Multiple studies have provided evidence that histone methylation is more important as a methyl sink than as a regulator of gene expression (Ye, Sutter, Wang, Kuang, & Tu, 2017; Ye et al., 2019). Under this hypothetical framework, *Kdm3* loss results in hypermethylation of histones. Hypermethylation may cause reduced recycling of methyl groups back into the folate cycle and reduced methyl group availability in the form of SAM. Indeed, HDM loss can decrease SAM (Ye et al., 2019). In response, other regulators of gene expression may upregulate 1-C genes to augment SAM output, leading to our observed expression changes. More work is needed to investigate these hypotheses.

**Fig. 7.**
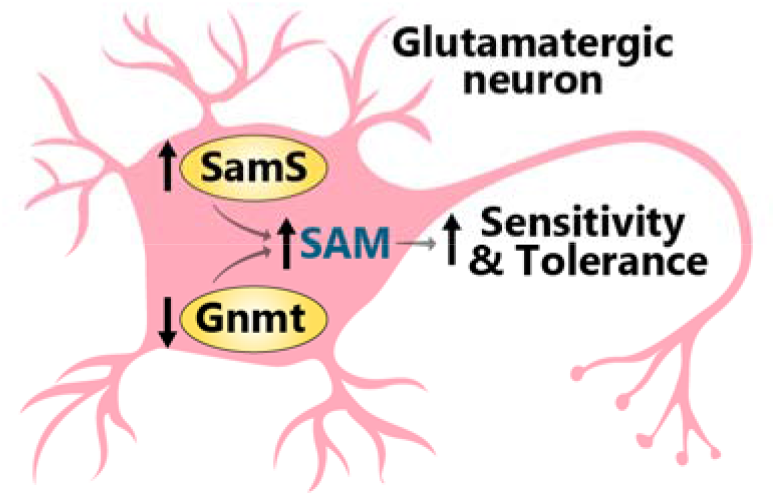
Increasing *S*-adenosyl methionine (SAM) in glutamatergic neurons enhances alcohol-induced sedation sensitivity and tolerance. Upregulating *SamS* or reducing *Gnmt* increases SAM levels in glutamatergic neurons, which in turn increases alcohol sensitivity and tolerance.

We also show that amino acid feedings, particularly of glycine, are sufficient to alter sensitivity and tolerance phenotypes. These effects disappear with *Kdm3* loss. The reason that casamino acids and glycine fail to affect *Kdm3*^*KO*^ flies is unknown but may be due to *Kdm3*^*KO*^-induced upregulation of *Gldc* and *Gnmt*, which buffer out excess glycine.

Global and neuronal 1-C manipulations induce changes in sensitivity and tolerance via different mechanisms. Globally, methyl group input into the folate cycle, as assessed via *Gldc* knockout and formate feeding, does not drive sensitivity and tolerance phenotypes. This may be true despite the neuronal importance of SAM because folate cycle inputs must pass several intermediate steps of regulation and buffering before those inputs can influence SAM. For instance, Wang et al. found that methionine depletion caused large drops in methylation levels, whereas serine or glycine loss caused only modest decreases (Wang et al., 2019). Leung et al. found that *Gldc* deficiency had no effect on the abundance of SAM, SAH, or the ratio of SAM/SAH (Leung et al., 2017). Further, mice with homozygous null mutation of a key folate cycle enzyme, *Mthfr*, exhibited no neural defects. Indeed, our results suggest that global glycine levels, rather than folate cycle methyl group levels, determine changes to alcohol sensitivity and tolerance, such that decreased sensitivity and tolerance are almost unanimously associated with expected global glycine elevation. These phenotypes are consistent even though we would expect some to increase SAM (e.g., *Gnmt* loss) and others to decrease SAM (e.g., cycloleucine feeding). Therefore, global glycine levels affect alcohol sensitivity and tolerance in a SAM-independent manner.

Further supporting this hypothesis, many studies have implicated glycine in AUD. EtOH targets and potentiates glycine receptors (Burgos, Muñoz, Guzman, & Aguayo, 2015; San Martin et al., 2020) and blocking the glial glycine transporter GlyT1 reduces EtOH consumption, preference, and relapse in rats (Molander, Lidö, Löf, Ericson, & Söderpalm, 2007; Vengeliene, Leonardi-Essmann, Sommer, Marston, & Spanagel, 2010). Similarly, systemic glycine treatment attenuates EtOH intake and preference in rats (Olsson, Höifödt Lidö, Danielsson, Ericson, & Söderpalm, 2021). In humans, fronto-cortical glycine levels are associated with recent heavy drinking (Prisciandaro et al., 2019).

Glycine may act by directly enhancing inhibitory glycinergic neurotransmission. The directionality of our results from neuronal *Gnmt* knockdowns and neuronal *Grd* or *GlyT* knockdowns does not support a glycinergic-dependent explanation of our observed neuronal phenotypes (i.e., both manipulations generally increased sensitivity and tolerance despite opposite expected effects on glycinergic signaling). However, inhibiting glycinergic signaling (via *Grd* and *GlyT* knockdowns) and feeding glycine or globally knocking down *Gnmt* changed alcohol sensitivity and tolerance in opposite directions, consistent with glycinergic neurotransmission mediating systemic glycine effects. Supporting this hypothesis, glycinergic signaling in rodents modulates alcohol sedation (Blednov, Benavidez, Homanics, & Harris, 2012; Quinlan, Ferguson, Jester, Firestone, & Homanics, 2002; San Martin et al., 2020), dopamine release (Lidö, Ericson, Marston, & Söderpalm, 2011; Molander, Löf, Stomberg, Ericson, & Söderpalm, 2005), and alcohol consumption (Molander et al., 2005; San Martin et al., 2020). Additionally, an EtOH-resistant knock-in mutation of GlyRs in mice increased alcohol consumption and EtOH-induced conditioned place preference (Muñoz et al., 2020). Importantly, it also substantially reduced EtOH sedation sensitivity (~40%) in a loss of righting reflex assay and reduced tolerance in a rotarod motor assay (Aguayo et al., 2014). These studies and our *Grd* and *GlyT* knockdowns suggest that glycine may affect alcohol sensitivity and tolerance via glycinergic activity.

Alternatively, glycine could act through its role as a required NMDAR co-agonist. NMDARs are a key locus of neural plasticity, a major target of EtOH inhibition, and a modulator of many EtOH phenotypes (Carpenter-Hyland & Chandler, 2006; Ron & Wang, 2009). Supporting this hypothesis, administration in rodents of an antagonist of the NMDAR glycine binding site limited alcohol-related reward learning (Biała & Kotlińska, 1999), dependence (Kotlińska, 2001), and withdrawal seizures (Kotlinska & Liljequist, 1996). Moreover, mutation of a fly NMDAR subunit altered EtOH sensitivity (B. Troutwine et al., 2019), and reduction of the same subunit reduced alcohol tolerance (Maiya et al., 2012). Multiple association studies have implicated NMDARs in human alcoholism (Karpyak, Geske, Colby, Mrazek, & Biernacka, 2012; J. H. Kim et al., 2006; Rujescu et al., 2005; Wernicke et al., 2003). Other studies suggest that glycine modulates and counteracts the inhibitory effect of EtOH on NMDARs (Buller, Larson, Morrisett, & Monaghan, 1995; Dildy-Mayfield & Leslie, 1991; Popp, Lickteig, & Lovinger, 1999; Rabe & Tabakoff, 1990; Woodward & Gonzales, 1990). Therefore, as NMDAR co-agonist, glycine can enhance glutamatergic excitatory tone. This outcome would reduce naïve sedation sensitivity, as we indeed observed. Glycine may also influence functional tolerance by potentiating NMDAR-mediated neuronal plasticity. Therefore, the role of glycine as NMDAR co-agonist may explain our results demonstrating that glycine levels modulate alcohol sensitivity and tolerance, though further research is needed to test this hypothesis.

We have furthermore shown distinct effects and mechanisms of whole-body and neuronal experiments. Global changes influence neuronal milieu, so we expect global *Gnmt* loss to increase both global glycine and neuronal SAM. Despite the activation of both mechanisms, however, global *Gnmt* loss recapitulates glycine feeding rather than neuronal *Gnmt* loss, indicating that the effects of augmented global glycine dominate over neuronal effects.

In contrast to global mechanisms, we find through *SamS* knockdown and overexpression that neuronal SAM levels (i.e., methylation potential) determine alcohol phenotypes, wherein higher SAM produces greater alcohol sensitivity and tolerance. Further supporting this hypothesis, *Gnmt* loss in flies raises SAM (Obata et al., 2014; Obata & Miura, 2015) and the SAM/SAH ratio (Obata et al., 2014), which is sometimes suggested as an alternate indicator of methylation potential. Our neuronal *Gnmt* knockdowns (i.e., SAM increases) increased sensitivity and tolerance. Thus, together with neuronal *SamS* knockdown and overexpression, we present three distinct lines of evidence consistently indicating that neuronal SAM increases alcohol sensitivity and tolerance. SAM is the universal donor of methyl groups for methyltransferase-mediated methylation reactions, including methylation of nucleic acids, lipids, histones, and other proteins. SAM is also critical to metabolic pathways such as synthesis of creatine, phosphatidylcholine, cysteine, and glutathione. Thus, SAM represents a powerful chokepoint capable of influencing metabolism, RNA processing, gene expression, protein translocation, signal transduction, and other protein and lipid functions. Intracellular SAM levels influence methylation rates, including histone methylation (Mentch & Locasale, 2016; Mentch et al., 2015; Shyh-Chang et al., 2013; Wang et al., 2019) (Liu et al., 2015; Liu & Pile, 2017), and even small fluctuations in SAM concentration may drastically alter HMT activity and methylation rates (Mentch & Locasale, 2016; Mentch et al., 2015). HDM loss enhances histone methylation (Liu et al., 2015; Liu & Pile, 2017), and neuronal loss of the HDM *Kdm3* increases EtOH sensitivity (Pinzon et al., 2017). Thus, these data support the hypothesis that greater methylation leads to greater sensitivity and tolerance.

In our study, pharmacologically or genetically reducing SAM yielded less alcohol sensitivity and tolerance. All our feedings were acute manipulations during the flies’ adulthood, suggesting that their effects on alcohol responses represent acute physiological changes, not developmental insults. It is unknown if such physiological changes arise from altered methylation in the brain or from homeostatic adaptations to changes in SAM levels. To shed light on this question, future studies should examine the impact of acutely elevated SAM on gene expression, histone methylation, and various metabolites associated with the 1-C cycles, both before and after ethanol exposure. One interesting possibility is that SAM levels affect alcohol phenotypes through SAM’s eventual conversion into glutathione, which reduces oxidative stress. SAM regulates glutathione levels (Ouyang, Wu, Li, Sun, & Sun, 2020). EtOH administration rapidly decreases glutathione (S. K. Kim, Seo, Jung, Kwak, & Kim, 2003), and glutathione and oxidative stress have previously been linked to the damaging effects of AUDs and to propensity to develop the disease (Björk et al., 2006; Covolo et al., 2005; Liang et al., 2004). Alternatively, protein methylation affects neurite outgrowth (Amano et al., 2020; Cimato, Ettinger, Zhou, & Aletta, 1997; Sontag, Nunbhakdi-Craig, Mitterhuber, Ogris, & Sontag, 2010), so elevated SAM may facilitate neuronal connections. These heightened connections may in turn contribute to faster spreading of EtOH-induced neuronal sedation and to greater neuronal plasticity, enhancing tolerance. Indeed, ethanol promotes growth of dendritic spines (Carpenter-Hyland & Chandler, 2006). Ultimately, future studies should better elucidate the mechanisms by which SAM modulates alcohol sensitivity and tolerance.

We have narrowed down SAM’s influence to glutamatergic neurons, while detecting no effects in inhibitory neurons. This contrast may suggest that *Drosophila* glutamatergic systems play a generally larger role in alcohol sensitivity and tolerance. Other studies have found that glutamatergic circuits modulate alcohol-associated memories (Scaplen et al., 2020). However, glutamate is not the primary excitatory neurotransmitter in the fly brain and thus may not be the best poised to alter excitatory/inhibitory homeostatic balance (Chvilicek, Titos, & Rothenfluh, 2020). Thus, glutamatergic neurons may not impact alcohol sensitivity and tolerance more than other neurons per se; instead, SAM may more powerfully influence sensitivity and tolerance via these neurons than it does via others, for reasons yet to be determined. Regardless, our glutamate data supports our previous hypothesis that glycine levels may influence alcohol sensitivity and tolerance by acting through glutamatergic NMDAR signaling. In fact, glutamatergic signaling could mediate all our observed phenotypes: higher glycine may enhance NMDAR activity to decrease sedation, while lower neuronal SAM could decrease methylation reactions, ultimately enhancing glutamatergic neurotransmission through unknown mechanisms and reducing sensitivity. Future studies should assess neuronal activity in various neuronal subpopulations and NMDAR activity as functions of glycine and SAM levels. Taken together, our study reveals a novel connection between epigenetic modifiers, 1-C metabolism, and alcohol responses that opens the door to greater understanding of AUDs.

## MATERIALS AND METHODS

### Fly Stocks and Husbandry

Flies were kept on standard cornmeal/agar medium at 25°C with 70% relative humidity. For all experiments, adult flies were sorted into separate vials 2-3 days prior to testing, and flies were generally 2-7 days post-eclosion at the time of experimentation. Male flies were used in all experiments except for those in Fig. 2 and Supp. 1 (because females consumed more of the special food) and *SamS* neuronal knockdown (which was lethal in males). *w\** Berlin flies were used as +/+ controls. *Kdm3*^*KO*^, *Gnmt*^*Mi*^, *Gldc*^*Mi*^, *elav*^*C155*^*-Gal4, tubulin84B-Gal4*, and *repo-Gal4* flies were outcrossed for at least five generations. In experiments using *Gnmt*^*RNAi 2*^ as the variable, the unexpressed RNAi construct was first confirmed to have no effect on alcohol responses. *Kdm3*^*KO*^ flies were generated in our previous study (Shalaby et al., 2017). Transgenic flies were obtained from the Bloomington Stock Center: *Gldc*^*Mi*^ (BDSC_59491), *Gnmt*^*Mi*^ (BDSC_67643), *Gnmt*^*RNAi 1*^ (BDSC_43148), *VGlut-Gal4* (BDSC_84697), *vgat3-Gal4* (BDSC_58409), *Shmt*^*KG*^ (BDSC_14948), and *Shmt*^*RNAi*^ (BDSC_57739). Dr. Clement Chow (University of Utah) provided the *repo-Gal4* flies. Dr. Carl Thummel (University of Utah) provided the *r4-Gal4* flies. Dr. Fernanda Ceriani (Fundación Instituto Leloir) graciously gifted us *Grd*^*RNAi*^ (VDRC v2702) and *GlyT*^*RNAi*^ (NIG-FLY Stocks, Transformant ID 5549R). Dr. Masayuki Miura and Dr. Fumiaki Obata (University of Tokyo) graciously gifted us *UAS-Gnmt*^*RNAi 2*^ (VDRC v25983), *UAS-Sardh*^*RNAi*^ (Obata et al., 2014; Obata & Miura, 2015), *UAS-SamS* (Obata & Miura, 2015), and *UAS-SamS*^*RNAi*^ (VDRC v103143).

### RNA-seq

Total RNA was isolated from heads of control and *Kdm3*^*KO*^ flies (50 flies per replicate; 3 replicates per genotype) using TRIzol Reagent (Invitrogen) and a PureLink RNA purification kit (ThermoFisher). Then, rRNA was removed from each sample with a Ribo-Zero rRNA Removal kit (Illumina). RNA libraries were constructed using a NEBNext Ultra II RNA Library Kit for Illumina and NEBNext Multiplex Oligos for Illumina, Primer Set 1 (New England Biolabs). Libraries were sequenced on an Illumina HiSeq 2500 instrument using 50-bp single-end reads.

### RNA-seq data analysis

Fastq files were aligned to the BDGP6 genome assembly using the STAR aligner (Dobin et al., 2013). The resulting BAM files were sorted and indexed using Samtools (Li et al., 2009). HTSeq count was used to collect count data (Anders, Pyl, & Huber, 2015). Differentially expressed genes were analyzed using the DEseq2 R package (Love, Huber, & Anders, 2014). Gene ontology analysis was performed using the ChIPseeker package in R (Yu, Wang, & He, 2015). Volcano plots were generated with the EnhancedVolcano R package (version 1.14.0, https://github.com/kevinblighe/EnhancedVolcano).

### Quantitative RT-PCR

Total RNA was purified from ~100 male *Kdm3*^*KO*^ fly heads using TRIzol Reagent (Invitrogen). RNAse-free glycogen was used as a carrier to precipitate isolated RNA. cDNA was made from 1 μg DNase-treated total RNA using iScript cDNA Synthesis Kits. Quantitative PCR was performed using Taqman Gene Expression Assays (ThermoFisher; Assay IDs: *Gnmt* – Dm02139745_g1; CG3999 – Dm02138658_g1; *Shmt* – Dm01796134_g1). Samples were run on an Applied Biosystems 7900HT real time PCR instrument using *RpL32* (ThermoFisher; Assay ID: Dm02151827_g1) as internal control.

### Specialized feedings

Adult flies were allowed to feed for three days ad libitum on food made from 1% (w/v) agar with 200 mM sucrose, with or without casamino acids (MP Biomedicals), glycine (Fisher Bioreagents), formate (Sigma), or cycloleucine (TCI Chemicals). Vials were replaced once if needed to prevent fungal/bacterial growth. Sensitivity and tolerance were assayed at the end of the three days.

### Ethanol sensitivity and tolerance assays

Maples assays were performed as previously described, with minor modifications (Maples & Rothenfluh, 2011). In brief, 10 flies were exposed to EtOH vapor for 22 minutes and scored for sedation. They were returned to their home vials at 25°C, then exposed again four hours after the start of first exposure to assess tolerance. Casamino acid feedings were performed using the Booze-o-mat as described previously (Wolf, Rodan, Tsai, & Heberlein, 2002). Briefly, flies were allowed to acclimate to air flow for 5 minutes. Then, the flies were exposed to 110/40 vaporized EtOH/air, which flowed for 31 minutes. Flies were considered sedated when they lost their self-righting ability. In all alcohol experiments, results were excluded for any vials in which ≥1/2 of the flies died.

### Blue Dye Consumption Assay

Flies were collected under CO_2_ anesthesia and allowed to recover for 24 hours in standard food vials. Flies were then transferred to vials with water-soaked cotton balls and strips of filter paper (7 × 1.75 cm) soaked with 350 ul feeding solution (200 mM sucrose, 0.68% (v/v) propionic acid, amino acids, and 0.3% (v/v) blue food dye (FD&C Blue Dye no. 1). After 24 hours, dead flies were removed and frozen at –20°C. Flies were grouped into sets of five and homogenized in 10 ul water. Homogenates were centrifuged for 5 minutes at 14,000 rpm. A NanoDrop spectrophotometer was used to measure 630 nm and 700 nm absorbance of the dyed supernatant, avoiding the topmost lipid layer. Feeding volume for each fly was calculated as *nL eaten = (OD 630 nm - 1*.*1*OD 700 nm)*CF*, where CF is a conversion factor specific to the Blue#1 stock solution.

### Data Analysis and Statistics

Statistical analyses were performed with GraphPad Prism 9. Previous studies indicate that sensitivity and tolerance phenotypes follow normal distributions (Pinzon et al., 2017), so we used Student’s t-tests for all two-group comparisons and standard 1-way ANOVA (for three-group comparisons) and 2-way ANOVA (for four-group comparisons), followed by Tukey’s post-hoc tests. Differences between standard deviations were tested with F tests and the Brown–Forsythe test. Where significant differences were found, Welch’s t-tests were used.

## ACKNOWLEDGMENTS

This work was supported by the University of Utah Genomics Core Facility, the High Throughput Sequencing Core at the Huntsman Cancer Institute, and the National Cancer Institute through Award Number 5P30CA042014. The content is solely the responsibility of the authors and does not necessarily represent the official views of the National Institutes of Health. We also thank all members of the Rothenfluh and Rodan labs for frequent feedback and support.

---

**Fig. S1.**
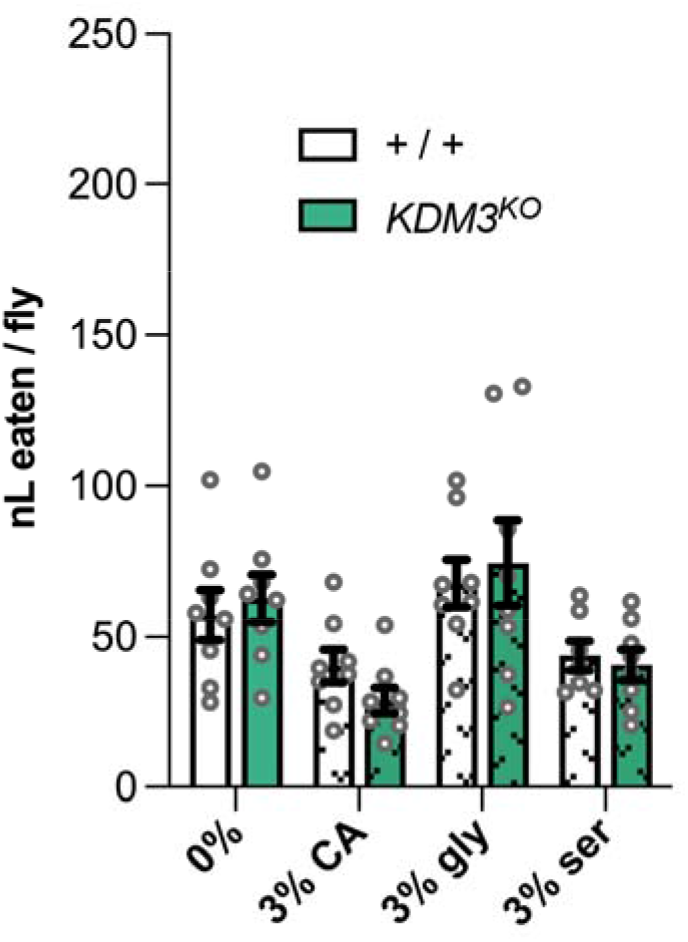
*Kdm3*^*KO*^ does not decrease amino acid consumption. Blue feeding analysis reveals that *Kdm3*^*KO*^ flies do not eat less amino acid-filled food than controls [CA, casamino acids; gly, glycine; ser, serine]. *n* = 9. Each dot represents five flies.

**Fig. S2.**
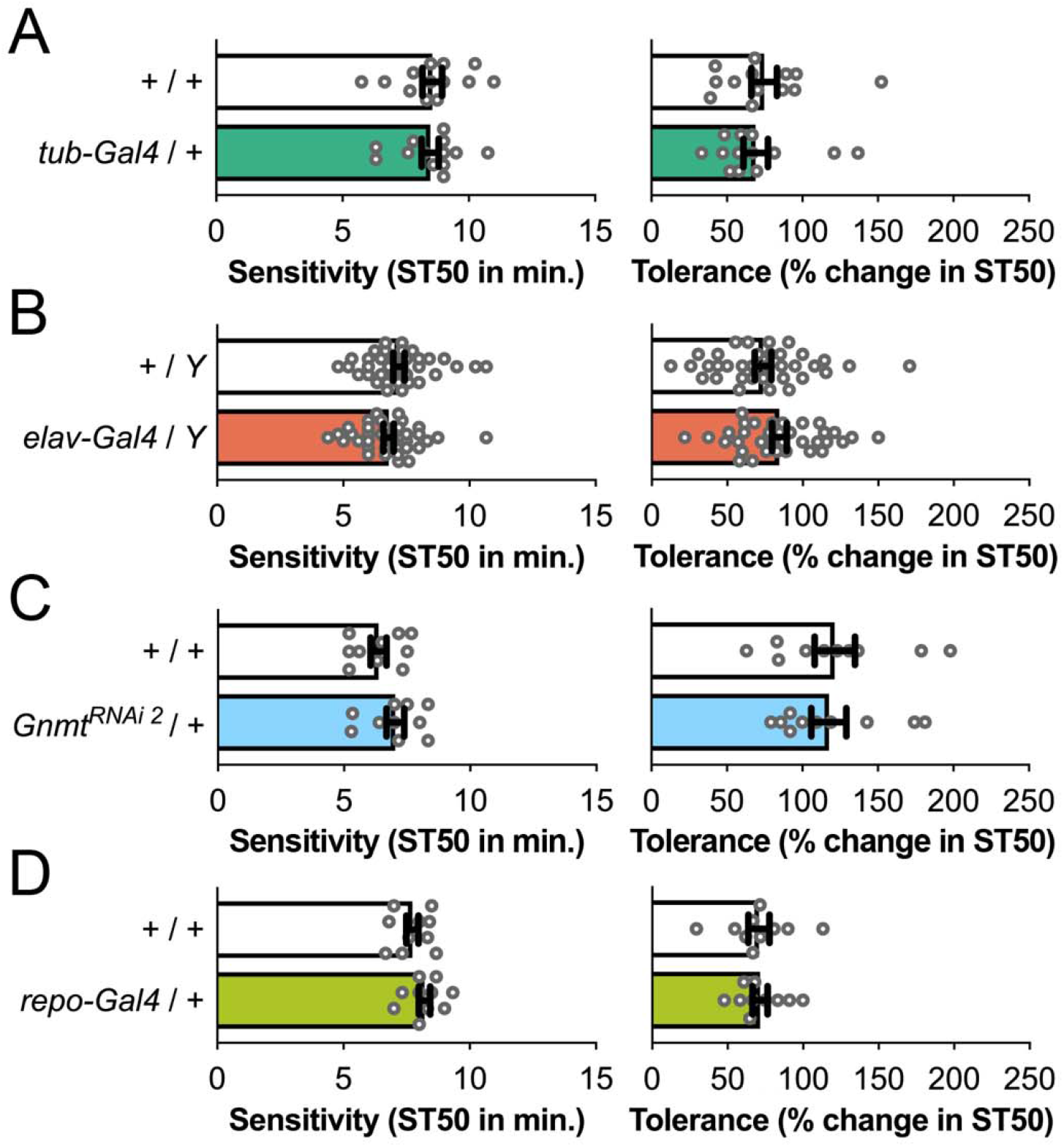
*Gal4* and UAS-RNAi constructs do not alter alcohol sensitivity and tolerance. (A) The *Tubulin84B-Gal4* global driver alone does not affect alcohol sensitivity or tolerance. (B) *elav-Gal4*, (C) *Gnmt*^*RNAi 2*^, nor (D) *repo-Gal4* affect alcohol sedation or tolerance. *n* ≥ 10.

**Fig. S3.**
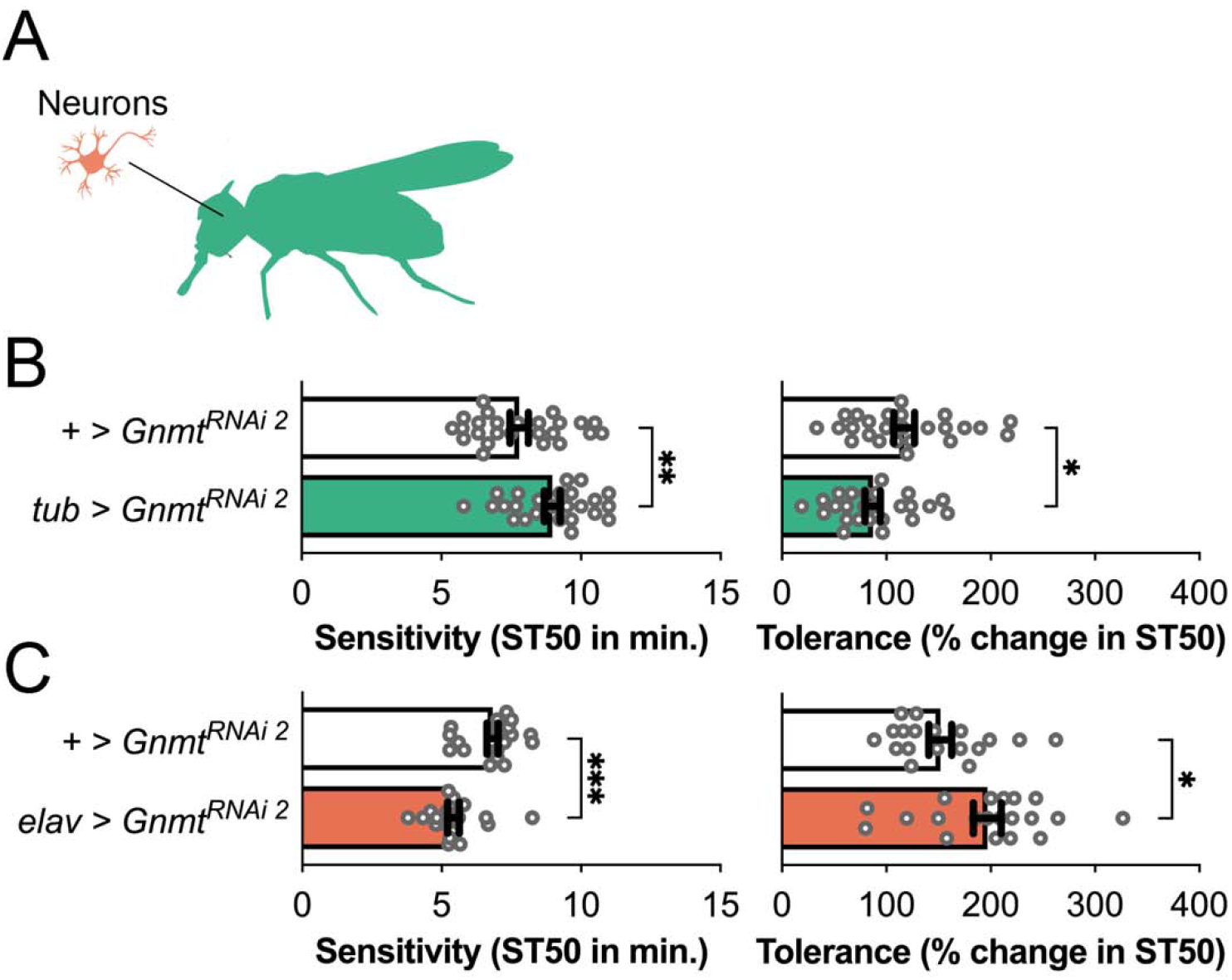
Second RNAi construct confirms global and neuronal *Gnmt* phenotypes. (A) Schematic of manipulation locations. Green represents whole-fly manipulations and orange represents neuronal manipulations. (B) Consistent with the first RNAi construct, global *Gnmt* knockdown using *Tubulin84B-Gal4* and *UAS-Gnmt*^*RNAi 2*^ decreases alcohol sensitivity and tolerance. (C) Neuronal *Gnmt* knockdown using *UAS-Gnmt*^*RNAi 2*^ increases alcohol sensitivity and tolerance, consistent with *UAS-Gnmt*^*RNAi 1*^. *n* ≥ 18.

**Fig. S4.**
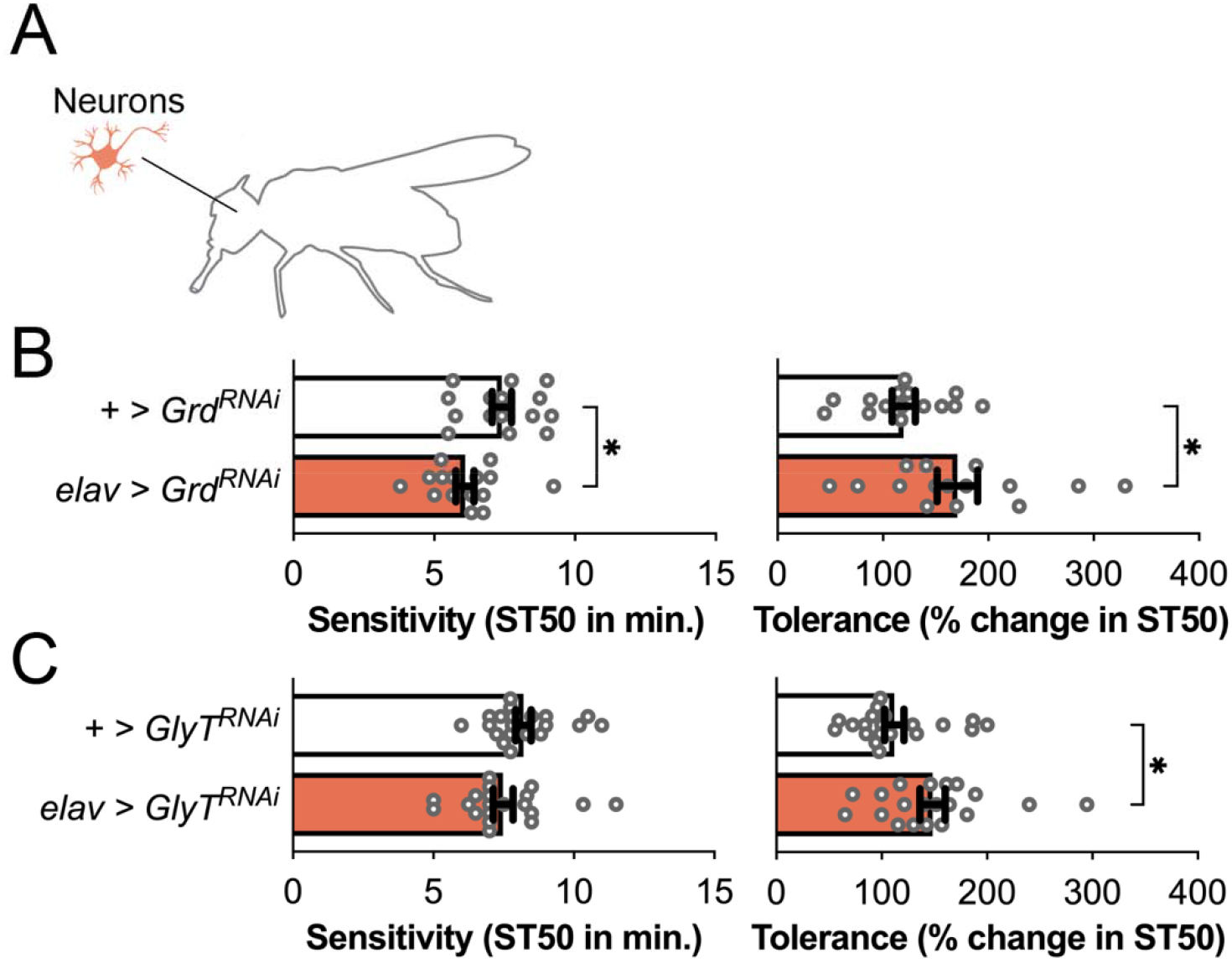
Glycinergic signaling plays a role in alcohol sensitivity and tolerance. (A) Schematic of experimental setup. (B) Pan-neuronal glycine receptor (*Grd*) knockdown increased alcohol sensitivity and tolerance. (C) Glycine reuptake transporter (*GlyT*) knockdown only increased tolerance. *n* ≥ 15.

**Supp. 5.**
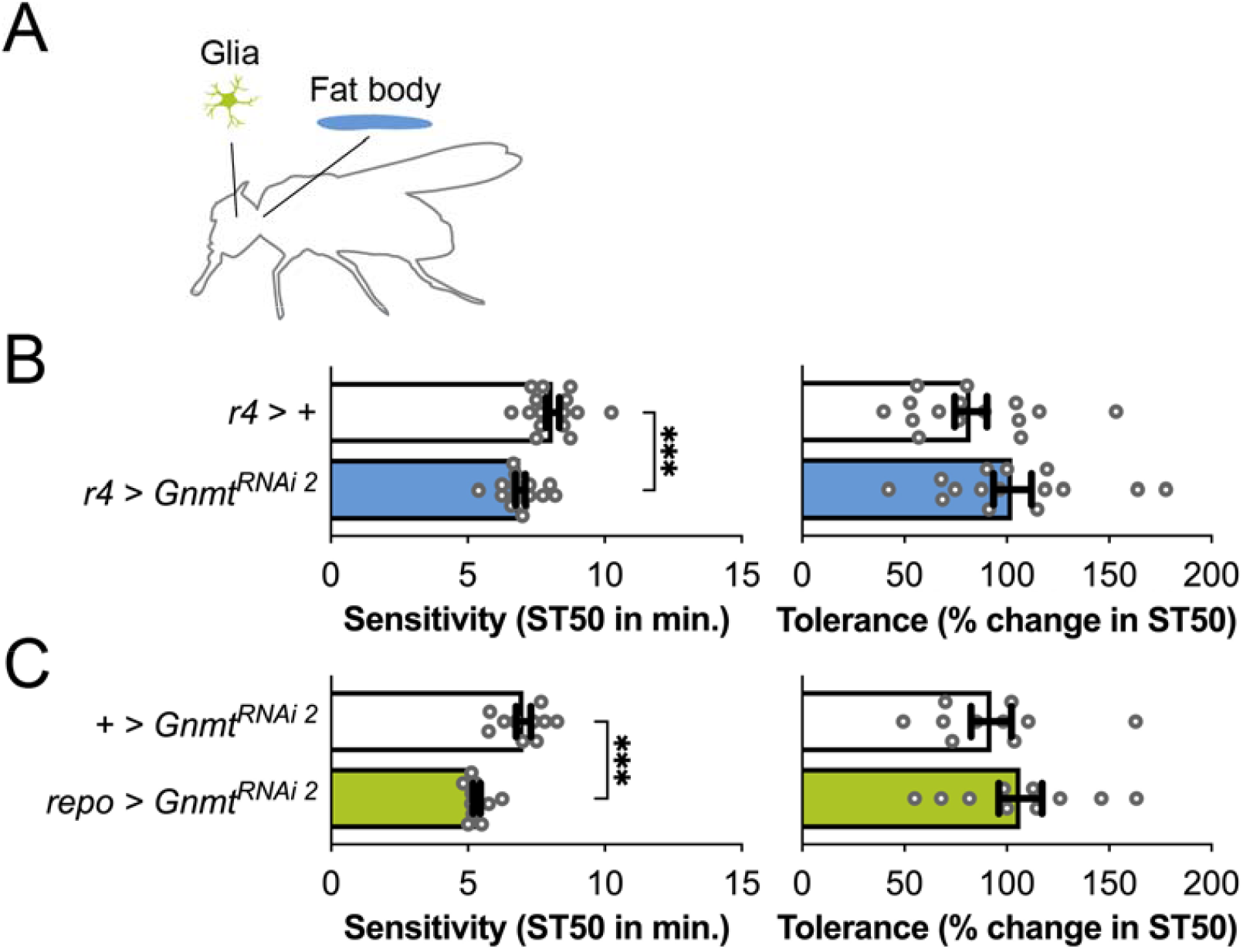
*Gnmt* knockdown in the fat body or glia recapitulates neuronal, but not global, *Gnmt* knockdown. (A) Color-coded schematic of manipulations in glia (olive) or the fat body (blue). (B) *Gnmt* knockdown in the fat body using *r4-Gal4*, a fat body-specific driver, increases sensitivity, consistent with neuronal phenotypes rather than global phenotypes. (C) *Gnmt* knockdown in glia using *repo-Gal4*, a glia-specific driver, causes the same result. *n* ≥ 10.

## REFERENCES

Aguayo, L. G., Castro, P., Mariqueo, T., Muñoz, B., Xiong, W., Zhang, L., … Homanics, G. E. (2014). Altered sedative effects of ethanol in mice with α1 glycine receptor subunits that are insensitive to Gβγ modulation. Neuropsychopharmacology, 39(11), 2538–2548. doi:10.1038/npp.2014.100

Amano, G., Matsuzaki, S., Mori, Y., Miyoshi, K., Han, S., Shikada, S., … Katayama, T. (2020). SCYL1 arginine methylation by PRMT1 is essential for neurite outgrowth via Golgi morphogenesis. Mol Biol Cell, 31(18), 1963–1973. doi:10.1091/mbc.E20-02-0100

Anders, S., Pyl, P. T., & Huber, W. (2015). HTSeq--a Python framework to work with high-throughput sequencing data. Bioinformatics, 31(2), 166–169. doi:10.1093/bioinformatics/btu638

Atkinson, N. S. (2009). Tolerance in Drosophila. J Neurogenet, 23(3), 293–302. doi:10.1080/01677060802572937

Aubrey, K. R., & Supplisson, S. (2018). Heterogeneous Signaling at GABA and Glycine Co-releasing Terminals. Front Synaptic Neurosci, 10, 40. doi:10.3389/fnsyn.2018.00040

Barbier, E., Johnstone, A. L., Khomtchouk, B. B., Tapocik, J. D., Pitcairn, C., Rehman, F., … Heilig, M. (2017). Dependence-induced increase of alcohol self-administration and compulsive drinking mediated by the histone methyltransferase PRDM2. Mol Psychiatry, 22(12), 1746–1758. doi:10.1038/mp.2016.131

Berger, K. H., Heberlein, U., & Moore, M. S. (2004). Rapid and chronic: two distinct forms of ethanol tolerance in Drosophila. Alcohol Clin Exp Res, 28(10), 1469–1480. doi:10.1097/01.alc.0000141817.15993.98

Berkel, T. D., & Pandey, S. C. (2017). Emerging Role of Epigenetic Mechanisms in Alcohol Addiction. Alcohol Clin Exp Res, 41(4), 666–680. doi:10.1111/acer.13338

Berkel, T. D. M., Zhang, H., Teppen, T., Sakharkar, A. J., & Pandey, S. C. (2019). Essential Role of Histone Methyltransferase G9a in Rapid Tolerance to the Anxiolytic Effects of Ethanol. Int J Neuropsychopharmacol, 22(4), 292–302. doi:10.1093/ijnp/pyy102

Biała, G., & Kotlińska, J. (1999). Blockade of the acquisition of ethanol-induced conditioned place preference by N-methyl-D-aspartate receptor antagonists. Alcohol Alcohol, 34(2), 175–182. doi:10.1093/alcalc/34.2.175

Björk, K., Saarikoski, S. T., Arlinde, C., Kovanen, L., Osei-Hyiaman, D., Ubaldi, M., … Sommer, W. H. (2006). Glutathione-S-transferase expression in the brain: possible role in ethanol preference and longevity. Faseb j, 20(11), 1826–1835. doi:10.1096/fj.06-5896com

Blednov, Y. A., Benavidez, J. M., Homanics, G. E., & Harris, R. A. (2012). Behavioral characterization of knockin mice with mutations M287L and Q266I in the glycine receptor α1 subunit. J Pharmacol Exp Ther, 340(2), 317–329. doi:10.1124/jpet.111.185124

Brosnan, M. E., & Brosnan, J. T. (2016). Formate: The Neglected Member of One-Carbon Metabolism. Annu Rev Nutr, 36, 369–388. doi:10.1146/annurev-nutr-071715-050738

Buller, A. L., Larson, H. C., Morrisett, R. A., & Monaghan, D. T. (1995). Glycine modulates ethanol inhibition of heteromeric N-methyl-D-aspartate receptors expressed in Xenopus oocytes. Mol Pharmacol, 48(4), 717–723.

Burgos, C. F., Muñoz, B., Guzman, L., & Aguayo, L. G. (2015). Ethanol effects on glycinergic transmission: From molecular pharmacology to behavior responses. Pharmacol Res, 101, 18–29. doi:10.1016/j.phrs.2015.07.002

Carpenter-Hyland, E. P., & Chandler, L. J. (2006). Homeostatic plasticity during alcohol exposure promotes enlargement of dendritic spines. Eur J Neurosci, 24(12), 3496–3506. doi:10.1111/j.1460-9568.2006.05247.x

Chvilicek, M. M., Titos, I., & Rothenfluh, A. (2020). The Neurotransmitters Involved in Drosophila Alcohol-Induced Behaviors. Front Behav Neurosci, 14, 607700. doi:10.3389/fnbeh.2020.607700

Cimato, T. R., Ettinger, M. J., Zhou, X., & Aletta, J. M. (1997). Nerve growth factor-specific regulation of protein methylation during neuronal differentiation of PC12 cells. J Cell Biol, 138(5), 1089–1103. doi:10.1083/jcb.138.5.1089

Covolo, L., Gelatti, U., Talamini, R., Garte, S., Trevisi, P., Franceschi, S., … Donato, F. (2005). Alcohol dehydrogenase 3, glutathione S-transferase M1 and T1 polymorphisms, alcohol consumption and hepatocellular carcinoma (Italy). Cancer Causes Control, 16(7), 831–838. doi:10.1007/s10552-005-2302-2

Dildy-Mayfield, J. E., & Leslie, S. W. (1991). Mechanism of inhibition of N-methyl-D-aspartate-stimulated increases in free intracellular Ca2+ concentration by ethanol. J Neurochem, 56(5), 1536–1543. doi:10.1111/j.1471-4159.1991.tb02048.x

Dobin, A., Davis, C. A., Schlesinger, F., Drenkow, J., Zaleski, C., Jha, S., … Gingeras, T. R. (2013). STAR: ultrafast universal RNA-seq aligner. Bioinformatics, 29(1), 15–21. doi:10.1093/bioinformatics/bts635

Engel, G. L., Marella, S., Kaun, K. R., Wu, J., Adhikari, P., Kong, E. C., & Wolf, F. W. (2016). Sir2/Sirt1 Links Acute Inebriation to Presynaptic Changes and the Development of Alcohol Tolerance, Preference, and Reward. J Neurosci, 36(19), 5241–5251. doi:10.1523/JNEUROSCI.0499-16.2016

Frenkel, L., Muraro, N. I., Beltran Gonzalez, A. N., Marcora, M. S., Bernabo, G., Hermann-Luibl, C., … Ceriani, M. F. (2017). Organization of Circadian Behavior Relies on Glycinergic Transmission. Cell Rep, 19(1), 72–85. doi:10.1016/j.celrep.2017.03.034

Friso, S., Udali, S., De Santis, D., & Choi, S. W. (2017). One-carbon metabolism and epigenetics. Mol Aspects Med, 54, 28–36. doi:10.1016/j.mam.2016.11.007

Ghezzi, A., Li, X., Lew, L. K., Wijesekera, T. P., & Atkinson, N. S. (2017). Alcohol-Induced Neuroadaptation Is Orchestrated by the Histone Acetyltransferase CBP. Front Mol Neurosci, 10, 103. doi:10.3389/fnmol.2017.00103

Goldman, D., Oroszi, G., & Ducci, F. (2005). The genetics of addictions: uncovering the genes. Nat Rev Genet, 6(7), 521–532. doi:10.1038/nrg1635

Gonzalez, D. A., Jia, T., Pinzon, J. H., Acevedo, S. F., Ojelade, S. A., Xu, B., … Rothenfluh, A. (2018). The Arf6 activator Efa6/PSD3 confers regional specificity and modulates ethanol consumption in Drosophila and humans. Mol Psychiatry, 23(3), 621–628. doi:10.1038/mp.2017.112

Kalant, H. (1998). Research on tolerance: what can we learn from history? Alcohol Clin Exp Res, 22(1), 67–76. doi:10.1111/j.1530-0277.1998.tb03618.x

Karpyak, V. M., Geske, J. R., Colby, C. L., Mrazek, D. A., & Biernacka, J. M. (2012). Genetic variability in the NMDA-dependent AMPA trafficking cascade is associated with alcohol dependence. Addict Biol, 17(4), 798–806. doi:10.1111/j.1369-1600.2011.00338.x

Kashio, S., Obata, F., Zhang, L., Katsuyama, T., Chihara, T., & Miura, M. (2016). Tissue nonautonomous effects of fat body methionine metabolism on imaginal disc repair in Drosophila. Proc Natl Acad Sci U S A, 113(7), 1835–1840. doi:10.1073/pnas.1523681113

Khanna, J. M., Kalant, H., Shah, G., & Weiner, J. (1991). Rapid tolerance as an index of chronic tolerance. Pharmacol Biochem Behav, 38(2), 427–432. doi:10.1016/0091-3057(91)90302-i

Kim, D., Fiske, B. P., Birsoy, K., Freinkman, E., Kami, K., Possemato, R. L., … Sabatini, D. M. (2015). SHMT2 drives glioma cell survival in ischaemia but imposes a dependence on glycine clearance. Nature, 520(7547), 363–367. doi:10.1038/nature14363

Kim, J. H., Park, M., Yang, S. Y., Jeong, B. S., Yoo, H. J., Kim, J.-W., … Kim, S. A. (2006). Association study of polymorphisms in N-methyl-d-aspartate receptor 2B subunits (GRIN2B) gene with Korean alcoholism. Neuroscience Research, 56(2), 220–223. doi:https://doi.org/10.1016/j.neures.2006.06.013

Kim, S. K., Seo, J. M., Jung, Y. S., Kwak, H. E., & Kim, Y. C. (2003). Alterations in hepatic metabolism of sulfur-containing amino acids induced by ethanol in rats. Amino Acids, 24(1-2), 103–110. doi:10.1007/s00726-002-0324-6

Kim, Y., Park, J., & Choi, Y. K. (2019). The Role of Astrocytes in the Central Nervous System Focused on BK Channel and Heme Oxygenase Metabolites: A Review. Antioxidants (Basel), 8(5). doi:10.3390/antiox8050121

Kotlińska, J. (2001). NMDA antagonists inhibit the development of ethanol dependence in rats. Pol J Pharmacol, 53(1), 47–50.

Kotlinska, J., & Liljequist, S. (1996). Oral administration of glycine and polyamine receptor antagonists blocks ethanol withdrawal seizures. Psychopharmacology (Berl), 127(3), 238-244. Retrieved from https://www.ncbi.nlm.nih.gov/pubmed/8912402

Lê, A. D., & Kiianmaa, K. (1988). Characteristics of ethanol tolerance in alcohol drinking (AA) and alcohol avoiding (ANA) rats. Psychopharmacology (Berl), 94(4), 479–483. doi:10.1007/bf00212841

Leung, K. Y., Pai, Y. J., Chen, Q., Santos, C., Calvani, E., Sudiwala, S., … Greene, N. D. E. (2017). Partitioning of One-Carbon Units in Folate and Methionine Metabolism Is Essential for Neural Tube Closure. Cell Rep, 21(7), 1795–1808. doi:10.1016/j.celrep.2017.10.072

Li, H., Handsaker, B., Wysoker, A., Fennell, T., Ruan, J., Homer, N., … Durbin, R. (2009). The Sequence Alignment/Map format and SAMtools. Bioinformatics, 25(16), 2078–2079. doi:10.1093/bioinformatics/btp352

Li, H., Janssens, J., De Waegeneer, M., Kolluru, S. S., Davie, K., Gardeux, V., … Zinzen, R. P. (2022). Fly Cell Atlas: A single-nucleus transcriptomic atlas of the adult fruit fly. Science, 375(6584), eabk2432. doi:10.1126/science.abk2432

Liang, T., Habegger, K., Spence, J. P., Foroud, T., Ellison, J. A., Lumeng, L., … Carr, L. G. (2004). Glutathione S-transferase 8-8 expression is lower in alcohol-preferring than in alcohol-nonpreferring rats. Alcohol Clin Exp Res, 28(11), 1622–1628. doi:10.1097/01.alc.0000145686.79141.57

Lidö, H. H., Ericson, M., Marston, H., & Söderpalm, B. (2011). A role for accumbal glycine receptors in modulation of dopamine release by the glycine transporter-1 inhibitor org25935. Front Psychiatry, 2, 8. doi:10.3389/fpsyt.2011.00008

Liu, M., Barnes, V. L., & Pile, L. A. (2015). Disruption of Methionine Metabolism in Drosophila melanogaster Impacts Histone Methylation and Results in Loss of Viability. G3 (Bethesda), 6(1), 121–132. doi:10.1534/g3.115.024273

Liu, M., & Pile, L. A. (2017). The Transcriptional Corepressor SIN3 Directly Regulates Genes Involved in Methionine Catabolism and Affects Histone Methylation, Linking Epigenetics and Metabolism. J Biol Chem, 292(5), 1970–1976. doi:10.1074/jbc.M116.749754

Love, M. I., Huber, W., & Anders, S. (2014). Moderated estimation of fold change and dispersion for RNA-seq data with DESeq2. Genome Biol, 15(12), 550. doi:10.1186/s13059-014-0550-8

Luka, Z., Capdevila, A., Mato, J. M., & Wagner, C. (2006). A glycine N-methyltransferase knockout mouse model for humans with deficiency of this enzyme. Transgenic Res, 15(3), 393–397. doi:10.1007/s11248-006-0008-1

Maiya, R., Lee, S., Berger, K. H., Kong, E. C., Slawson, J. B., Griffith, L. C., … Heberlein, U. (2012). DlgS97/SAP97, a neuronal isoform of discs large, regulates ethanol tolerance. PLoS One, 7(11), e48967. doi:10.1371/journal.pone.0048967

Maples, T., & Rothenfluh, A. (2011). A simple way to measure ethanol sensitivity in flies. J Vis Exp(48), 2541. doi:10.3791/2541

Mayfield, R. D., Harris, R. A., & Schuckit, M. A. (2008). Genetic factors influencing alcohol dependence. Br J Pharmacol, 154(2), 275–287. doi:10.1038/bjp.2008.88

Maze, I., Covington, H. E., 3rd, Dietz, D. M., LaPlant, Q., Renthal, W., Russo, S. J., … Nestler, E. J. (2010). Essential role of the histone methyltransferase G9a in cocaine-induced plasticity. Science, 327(5962), 213–216. doi:10.1126/science.1179438

Mentch, S. J., & Locasale, J. W. (2016). One-carbon metabolism and epigenetics: understanding the specificity. Ann N Y Acad Sci, 1363, 91–98. doi:10.1111/nyas.12956

Mentch, S. J., Mehrmohamadi, M., Huang, L., Liu, X., Gupta, D., Mattocks, D., … Locasale, J. W. (2015). Histone Methylation Dynamics and Gene Regulation Occur through the Sensing of One-Carbon Metabolism. Cell metabolism, 22(5), 861–873. doi:10.1016/j.cmet.2015.08.024

Molander, A., Lidö, H. H., Löf, E., Ericson, M., & Söderpalm, B. (2007). The glycine reuptake inhibitor Org 25935 decreases ethanol intake and preference in male wistar rats. Alcohol Alcohol, 42(1), 11–18. doi:10.1093/alcalc/agl085

Molander, A., Löf, E., Stomberg, R., Ericson, M., & Söderpalm, B. (2005). Involvement of accumbal glycine receptors in the regulation of voluntary ethanol intake in the rat. Alcohol Clin Exp Res, 29(1), 38–45. doi:10.1097/01.alc.0000150009.78622.e0

Mulligan, M. K., Ponomarev, I., Hitzemann, R. J., Belknap, J. K., Tabakoff, B., Harris, R. A., … Bergeson, S. E. (2006). Toward understanding the genetics of alcohol drinking through transcriptome meta-analysis. Proc Natl Acad Sci U S A, 103(16), 6368–6373. doi:10.1073/pnas.0510188103

Muñoz, B., Gallegos, S., Peters, C., Murath, P., Lovinger, D. M., Homanics, G. E., & Aguayo, L. G. (2020). Influence of nonsynaptic α1 glycine receptors on ethanol consumption and place preference. Addict Biol, 25(2), e12726. doi:10.1111/adb.12726

Narayanan, A. S., & Rothenfluh, A. (2016). I Believe I Can Fly!: Use of Drosophila as a Model Organism in Neuropsychopharmacology Research. Neuropsychopharmacology, 41(6), 1439–1446. doi:10.1038/npp.2015.322

Obata, F., Kuranaga, E., Tomioka, K., Ming, M., Takeishi, A., Chen, C. H., … Miura, M. (2014). Necrosis-driven systemic immune response alters SAM metabolism through the FOXO-GNMT axis. Cell Rep, 7(3), 821–833. doi:10.1016/j.celrep.2014.03.046

Obata, F., & Miura, M. (2015). Enhancing S-adenosyl-methionine catabolism extends Drosophila lifespan. Nat Commun, 6, 8332. doi:10.1038/ncomms9332

Ojelade, S. A., Acevedo, S. F., Kalahasti, G., Rodan, A. R., & Rothenfluh, A. (2015). RhoGAP18B Isoforms Act on Distinct Rho-Family GTPases and Regulate Behavioral Responses to Alcohol via Cofilin. PLoS One, 10(9), e0137465. doi:10.1371/journal.pone.0137465

Ojelade, S. A., Jia, T., Rodan, A. R., Chenyang, T., Kadrmas, J. L., Cattrell, A., … Rothenfluh, A. (2015). Rsu1 regulates ethanol consumption in Drosophila and humans. Proc Natl Acad Sci U S A, 112(30), E4085–4093. doi:10.1073/pnas.1417222112

Olsson, Y., Höifödt Lidö, H., Danielsson, K., Ericson, M., & Söderpalm, B. (2021). Effects of systemic glycine on accumbal glycine and dopamine levels and ethanol intake in male Wistar rats. J Neural Transm (Vienna), 128(1), 83–94. doi:10.1007/s00702-020-02284-x

Ouyang, Y., Wu, Q., Li, J., Sun, S., & Sun, S. (2020). S-adenosylmethionine: A metabolite critical to the regulation of autophagy. Cell Prolif, 53(11), e12891. doi:10.1111/cpr.12891

Pai, Y. J., Leung, K. Y., Savery, D., Hutchin, T., Prunty, H., Heales, S., … Greene, N. D. (2015). Glycine decarboxylase deficiency causes neural tube defects and features of non-ketotic hyperglycinemia in mice. Nat Commun, 6, 6388. doi:10.1038/ncomms7388

Park, A., Ghezzi, A., Wijesekera, T. P., & Atkinson, N. S. (2017). Genetics and genomics of alcohol responses in Drosophila. Neuropharmacology, 122, 22–35. doi:10.1016/j.neuropharm.2017.01.032

Pinzon, J. H., Reed, A. R., Shalaby, N. A., Buszczak, M., Rodan, A. R., & Rothenfluh, A. (2017). Alcohol-Induced Behaviors Require a Subset of Drosophila JmjC-Domain Histone Demethylases in the Nervous System. Alcohol Clin Exp Res, 41(12), 2015–2024. doi:10.1111/acer.13508

Ponomarev, I. (2013). Epigenetic control of gene expression in the alcoholic brain. Alcohol Res, 35(1), 69–76. Retrieved from https://www.ncbi.nlm.nih.gov/pubmed/24313166

Ponomarev, I., Wang, S., Zhang, L., Harris, R. A., & Mayfield, R. D. (2012). Gene coexpression networks in human brain identify epigenetic modifications in alcohol dependence. J Neurosci, 32(5), 1884–1897. doi:10.1523/JNEUROSCI.3136-11.2012

Popp, R. L., Lickteig, R. L., & Lovinger, D. M. (1999). Factors that enhance ethanol inhibition of N-methyl-D-aspartate receptors in cerebellar granule cells. J Pharmacol Exp Ther, 289(3), 1564–1574.

Prisciandaro, J. J., Schacht, J. P., Prescot, A. P., Brenner, H. M., Renshaw, P. F., Brown, T. R., & Anton, R. F. (2019). Evidence for a unique association between fronto-cortical glycine levels and recent heavy drinking in treatment naïve individuals with alcohol use disorder. Neurosci Lett, 706, 207–210. doi:10.1016/j.neulet.2019.05.030

Qiang, M., Denny, A., Lieu, M., Carreon, S., & Li, J. (2011). Histone H3K9 modifications are a local chromatin event involved in ethanol-induced neuroadaptation of the NR2B gene. Epigenetics, 6(9), 1095–1104. doi:10.4161/epi.6.9.16924

Quinlan, J. J., Ferguson, C., Jester, K., Firestone, L. L., & Homanics, G. E. (2002). Mice with glycine receptor subunit mutations are both sensitive and resistant to volatile anesthetics. Anesth Analg, 95(3), 578–582, table of contents. doi:10.1097/00000539-200209000-00016

Rabe, C. S., & Tabakoff, B. (1990). Glycine site-directed agonists reverse the actions of ethanol at the N-methyl-D-aspartate receptor. Mol Pharmacol, 38(6), 753–757.

Ramirez-Roman, M. E., Billini, C. E., & Ghezzi, A. (2018). Epigenetic Mechanisms of Alcohol Neuroadaptation: Insights from Drosophila. J Exp Neurosci, 12, 1179069518779809. doi:10.1177/1179069518779809

Rizki, T. M., & Rizki, R. M. (1978). Larval adipose tissue of homoeotic bithorax mutants of Drosophila. Dev Biol, 65(2), 476–482. doi:10.1016/0012-1606(78)90042-8

Robinson, B. G., & Atkinson, N. S. (2013). Is alcoholism learned? Insights from the fruit fly. Curr Opin Neurobiol, 23(4), 529–534. doi:10.1016/j.conb.2013.01.016

Rodan, A. R., Kiger, J. A., Jr., & Heberlein, U. (2002). Functional dissection of neuroanatomical loci regulating ethanol sensitivity in Drosophila. J Neurosci, 22(21), 9490–9501. Retrieved from https://www.ncbi.nlm.nih.gov/pubmed/12417673

Rodan, A. R., & Rothenfluh, A. (2010). The genetics of behavioral alcohol responses in Drosophila. Int Rev Neurobiol, 91, 25–51. doi:10.1016/S0074-7742(10)91002-7

Ron, D., & Wang, J. (2009). The NMDA Receptor and Alcohol Addiction. In A. M. Van Dongen (Ed.), Biology of the NMDA Receptor. Boca Raton (FL): CRC Press/Taylor & Francis Taylor & Francis Group, LLC.

Rujescu, D., Soyka, M., Dahmen, N., Preuss, U., Hartmann, A. M., Giegling, I., … Szegedi, A. (2005). GRIN1 locus may modify the susceptibility to seizures during alcohol withdrawal. Am J Med Genet B Neuropsychiatr Genet, 133b(1), 85–87. doi:10.1002/ajmg.b.30112

Rustay, N. R., & Crabbe, J. C. (2004). Genetic analysis of rapid tolerance to ethanol’s incoordinating effects in mice: inbred strains and artificial selection. Behav Genet, 34(4), 441–451. doi:10.1023/B:BEGE.0000023649.60539.dd

San Martin, L., Gallegos, S., Araya, A., Romero, N., Morelli, G., Comhair, J., … Aguayo, L. G. (2020). Ethanol consumption and sedation are altered in mice lacking the glycine receptor α2 subunit. Br J Pharmacol, 177(17), 3941–3956. doi:10.1111/bph.15136

Scaplen, K. M., Talay, M., Nunez, K. M., Salamon, S., Waterman, A. G., Gang, S., … Kaun, K. R. (2020). Circuits that encode and guide alcohol-associated preference. Elife, 9. doi:10.7554/eLife.48730

Scholz, H., Ramond, J., Singh, C. M., & Heberlein, U. (2000). Functional ethanol tolerance in Drosophila. Neuron, 28(1), 261–271. doi:10.1016/s0896-6273(00)00101-x

Schuckit, M. A. (2009). An overview of genetic influences in alcoholism. J Subst Abuse Treat, 36(1), S5–14. Retrieved from https://www.ncbi.nlm.nih.gov/pubmed/19062348

Serefidou, M., Venkatasubramani, A. V., & Imhof, A. (2019). The Impact of One Carbon Metabolism on Histone Methylation. Front Genet, 10, 764. doi:10.3389/fgene.2019.00764

Shalaby, N. A., Sayed, R., Zhang, Q., Scoggin, S., Eliazer, S., Rothenfluh, A., & Buszczak, M. (2017). Systematic discovery of genetic modulation by Jumonji histone demethylases in Drosophila. Sci Rep, 7(1), 5240. doi:10.1038/s41598-017-05004-w

Shukla, S. D., Velazquez, J., French, S. W., Lu, S. C., Ticku, M. K., & Zakhari, S. (2008). Emerging role of epigenetics in the actions of alcohol. Alcohol Clin Exp Res, 32(9), 1525–1534. doi:10.1111/j.1530-0277.2008.00729.x

Shyh-Chang, N., Locasale, J. W., Lyssiotis, C. A., Zheng, Y., Teo, R. Y., Ratanasirintrawoot, S., … Cantley, L. C. (2013). Influence of threonine metabolism on S-adenosylmethionine and histone methylation. Science, 339(6116), 222–226. doi:10.1126/science.1226603

Sontag, J. M., Nunbhakdi-Craig, V., Mitterhuber, M., Ogris, E., & Sontag, E. (2010). Regulation of protein phosphatase 2A methylation by LCMT1 and PME-1 plays a critical role in differentiation of neuroblastoma cells. J Neurochem, 115(6), 1455–1465. doi:10.1111/j.1471-4159.2010.07049.x

Subbanna, S., Shivakumar, M., Umapathy, N. S., Saito, M., Mohan, P. S., Kumar, A., … Basavarajappa, B. S. (2013). G9a-mediated histone methylation regulates ethanol-induced neurodegeneration in the neonatal mouse brain. Neurobiology of Disease, 54, 475–485. doi:10.1016/j.nbd.2013.01.022

Sufrin, J. R., Coulter, A. W., & Talalay, P. (1979). Structural and conformational analogues of L-methionine as inhibitors of the enzymatic synthesis of S-adenosyl-L-methionine. IV. Further mono-, bi- and tricyclic amino acids. Mol Pharmacol, 15(3), 661–677.

Troutwine, B., Park, A., Velez-Hernandez, M. E., Lew, L., Mihic, S. J., & Atkinson, N. S. (2019). F654A and K558Q Mutations in NMDA Receptor 1 Affect Ethanol-Induced Behaviors in Drosophila. Alcohol Clin Exp Res, 43(12), 2480–2493. doi:10.1111/acer.14215

Troutwine, B. R., Ghezzi, A., Pietrzykowski, A. Z., & Atkinson, N. S. (2016). Alcohol resistance in Drosophila is modulated by the Toll innate immune pathway. Genes Brain Behav, 15(4), 382–394. doi:10.1111/gbb.12288

Vengeliene, V., Leonardi-Essmann, F., Sommer, W. H., Marston, H. M., & Spanagel, R. (2010). Glycine transporter-1 blockade leads to persistently reduced relapse-like alcohol drinking in rats. Biol Psychiatry, 68(8), 704–711. doi:10.1016/j.biopsych.2010.05.029

Venken, K. J., Schulze, K. L., Haelterman, N. A., Pan, H., He, Y., Evans-Holm, M., … Bellen, H. J. (2011). MiMIC: a highly versatile transposon insertion resource for engineering Drosophila melanogaster genes. Nature methods, 8(9), 737–743. doi:10.1038/nmeth.1662

Wang, Z., Yip, L. Y., Lee, J. H. J., Wu, Z., Chew, H. Y., Chong, P. K. W., … Tam, W. L. (2019). Methionine is a metabolic dependency of tumor-initiating cells. Nature Medicine, 25(5), 825–837. doi:10.1038/s41591-019-0423-5

Wernicke, C., Samochowiec, J., Schmidt, L. G., Winterer, G., Smolka, M., Kucharska-Mazur, J., … Rommelspacher, H. (2003). Polymorphisms in the N-methyl-D-aspartate receptor 1 and 2B subunits are associated with alcoholism-related traits. Biol Psychiatry, 54(9), 922–928. doi:10.1016/s0006-3223(03)00072-6

Wolf, F. W., Rodan, A. R., Tsai, L. T., & Heberlein, U. (2002). High-resolution analysis of ethanol-induced locomotor stimulation in Drosophila. J Neurosci, 22(24), 11035–11044. doi:10.1523/jneurosci.22-24-11035.2002

Woodward, J. J., & Gonzales, R. A. (1990). Ethanol inhibition of N-methyl-D-aspartate-stimulated endogenous dopamine release from rat striatal slices: reversal by glycine. J Neurochem, 54(2), 712–715. doi:10.1111/j.1471-4159.1990.tb01931.x

Ye, C., Sutter, B. M., Wang, Y., Kuang, Z., & Tu, B. P. (2017). A Metabolic Function for Phospholipid and Histone Methylation. Mol Cell, 66(2), 180-193.e188. doi:10.1016/j.molcel.2017.02.026

Ye, C., Sutter, B. M., Wang, Y., Kuang, Z., Zhao, X., Yu, Y., & Tu, B. P. (2019). Demethylation of the Protein Phosphatase PP2A Promotes Demethylation of Histones to Enable Their Function as a Methyl Group Sink. Mol Cell, 73(6), 1115–1126 e1116. doi:10.1016/j.molcel.2019.01.012

Yu, G., Wang, L. G., & He, Q. Y. (2015). ChIPseeker: an R/Bioconductor package for ChIP peak annotation, comparison and visualization. Bioinformatics, 31(14), 2382–2383. doi:10.1093/bioinformatics/btv145

Zakhari, S. (2013). Alcohol metabolism and epigenetics changes. Alcohol Res, 35(1), 6–16. Retrieved from https://www.ncbi.nlm.nih.gov/pubmed/24313160

Zinke, I., Kirchner, C., Chao, L. C., Tetzlaff, M. T., & Pankratz, M. J. (1999). Suppression of food intake and growth by amino acids in Drosophila: the role of pumpless, a fat body expressed gene with homology to vertebrate glycine cleavage system. Development, 126(23), 5275–5284. doi:10.1242/dev.126.23.5275

